# neoDL: A novel neoantigen intrinsic feature-based deep learning model identifies IDH wild-type glioblastomas with the longest survival

**DOI:** 10.1101/2020.12.28.424562

**Authors:** Ting Sun, Yufei He, Wendong Li, Guang Liu, Lin Li, Lu Wang, Zixuan Xiao, Xiaohan Han, Hao Wen, Yong Liu, Yifan Chen, Haoyu Wang, Jing Li, Yubo Fan, Wei Zhang, Jing Zhang

## Abstract

**Background:** IDH wild-type glioblastoma (GBM) is the most aggressive tumor in the central nervous system in spite of extensive therapies. Neoantigen based personalized immune therapies achieve promising results in melanoma and lung cancer, but few neoantigen based models perform well in IDH wild-type GBM. Unlike the neoantigen load and occurrence that are well studied and often found useless, the association between neoantigen intrinsic features and prognosis remain unclear in IDH wild-type GBM.

**Results:** We presented a novel neoantigen intrinsic feature-based deep learning model (neoDL) to stratify IDH wild-type GBMs into subgroups with different survivals. We first calculated a total of 2928 intrinsic features for each neoantigen and filtered out those not associated with survival, followed by applying neoDL in the TCGA data cohort. Leave one out cross validation (LOOCV) in the TCGA demonstrated that neoDL successfully classified IDH wild-type GBMs into different prognostic subgroups, which was further validated in an independent data cohorts from Asian population. Long-term survival IDH wild-type GBMs identified by neoDL were found characterized by 12 protective neoantigen intrinsic features and enriched in development and cell cycle.

**Conclusions:** Our results provide a novel model, neoDL, that can be therapeutically exploited to identify IDH wild-type GBM with good prognosis who will most likely benefit from neoantigen based personalized immunetherapy.

## Background

Glioblastoma is the most common aggressive primary brain tumor associated with profound genomic heterogeneity and high recurrence rate, which has limited therapy development[1–3]. Currently, the standard therapy for GBM is surgical resection, followed by radiotherapy and adjuvant chemotherapy[4, 5]. Although the survival status of GBM patients has improved with the advancement of modern combination therapies, the overall prognosis of the majority of GBM patients remains poor and clinical outcomes vary considerably among patients[6]. GBM still has the worst 5-year overall survival rates among all human cancers[7], with a dismal median duration of 14 months[8, 9].

Tumor neoantigens, classically regarded as being derived from mutation-containing proteins that generate novel immunogenic epitopes[10]. High nonsynonymous coding mutational loads harbor more neoantigens that are presented to CD8+ T cells on restricted HLA-I subtypes[11–13], leading to stronger immunogenicity and better overall survival in selected tumor types such as melanoma[14], lung cancer[15], and colorectal tumors[16]. However, in gliomas, higher mutational load means increased tumor aggressiveness[17]. Neoantigens are the attractive targets in personalized immunetherapies, promoting tumor-specific T-cell responses and affecting antitumor immune responses in a number of preclinical models[18, 19]. Additionally, optimized high-quality neoantigens model has been applied recently for the identification of GBM patients, which performed optimally in providing a more accurate prediction of anti-tumor immunity and identified patients with the longest survival[20]. Although the occurrence and characterization of neoantigen in pan-cancer has been studied, showing that all positions in neoepitopes of all lengths containing more hydrophobic residues than the wild-type sequences[21], the comprehensive features of neoantigens associated with prognosis and immunoreaction in IDH wild-type GBM remain elusive.

Deep learning models data by learning high-level representations and could learn features from noisy and raw data[22, 23]. It has outperformed traditional machine learning methods, especially in image and natural language processing[24–26]. The remarkable flexibility and adaptability of deep learning models have led to the proliferation of their application in bioinformatics research, such as protein structure prediction, biomedical imaging, and biomedical signal processing[26]. In recent years, image-based deep learning model has been widely used in cancer study and has shown excellent level-accuracy in precise diagnosis and prognostic stratification, such as colorectal[27], prostate[28, 29], melanoma[30], and gliomas[31]. Deep learning has also demonstrated its strong ability in prediction. Glioma grading[32], glioma genetic mutation[33] or survival[34] have all be predicted in previous researches. Recently, it also demonstrated that the application of neoantigen-based machine learning could successfully predict neoantigen immunogenicity in colon and lung adenocarcinomas [35].

In this manuscript, we present a deep learning model, which was designed based on the intrinsic features of neoantigens and could successfully stratify IDH wild-type GBM patients of TCGA into subgroups with different survival. This model was further validated in an independent data from Asian population, demonstrating its powerful predictive effects in some higher-grade glioma subtypes, including Classical, Classical-like, Glioblastoma, IDH wild-type, Mesenchymal-like. The features-based deep learning model informed a cohort of patients with better clinical prognosis, which generally exhibits biological processes such as development, and cell cycle. Our model for IDH wild-type GBM has important implications in diagnosis and prognosis, and also helps in identify patients who most likely benefit from neoantigen based personalized immunetherapy.

## Data Description

Mutations and clinical information were downloaded from the ATLAS-TCGA pan-glioma study[36]. Gene expression microarray data with Agilent chip (G4502A) at level 3 were downloaded from TCGA Data portal. We termed the data from TCGA as TCGA cohort. Mutations, RNAseq gene expression data, and clinical information in Asian population were collected from a recently published cohort[37], designated as Pri cohort. The samples that were not diagnosed as IDH wild-type GBM or did not have clinical information were removed, resulting in 268 and 46 samples in the two cohorts, respectively.

A neoepitope with strong affinity for MHC (*IC*_50_ equal or less than 500 nM) may be a more robust neoantigen candidate if the paired wild-type epitope has a poor affinity for MHC (*IC*_50_ greater than 500 nM)[38]. The neoantigens for each sample in both TCGA cohort and Pri cohort were from our previous study[20], which used missense mutations to generate all possible 9-mer peptides and defined the mutant 9-mer peptides as neoantigens when the *IC*_50_ of mutant-type 9-mer peptides was less than 500 nM and the corresponding wild-type binder more than 500 nM. All the downstream analyses were based on the inferred neoantigens (the mutant peptides) and their corresponding wild-type peptides.

## Analyses

### Identification of sequence features of neoantigens associated with the overall survival of IDH wild-type GBMs

Tumor mutational burden has been described as a predictor of tumor behavior and immunological response[39–41]. Recent researches showed that tumor mutational load correlated with improved survival and immunotherapy response in some tumors, including melanomas[42], ovarian[39], and bladder carcinoma[43]. To estimate the relationship between missense mutational load and overall survival, we calculated missense mutational load for 262 and 42 IDH wild-type GBMs in TCGA and Pri-cohort, respectively. Kaplan-Meier analysis demonstrated that there were no statistically significantly different overall survival between higher and lower mutation loads of IDH wild-type GBMs in both TCGA cohort and the Pri cohort (Figure 1A-B), consistent with the previous research[20]. Similarly, patients in different glioma subgroups containing high (above mean) or low (below mean) mutational loads had similar prognosis with Astrocytoma, Codel, OligoAstrocytoma, IDH MT-codel, IDH MT-noncodel having higher mutation loads related to worse clinical outcome (Supplementary Figure S1). High missense mutational load was predicted to harbor more short peptides including the mutation, leading to more neoantigens, rendering them more susceptible T-cell targets[44]. The absolute number of neoantigens, which was calculated by adding up all the mutant peptides for one sample, also failed to predict the survival of IDH wild-type GBMs (Figure 1C-D) and different glioma subgroups (Supplementary Figure S2). DAI, defined as difference between binding affinity of wildtype and mutant-type peptides for MHC class I, was reported to be a better indicator of immunogenicity than mutant affinity and a predictor of survival in advanced lung cancer and melanoma[45]. We calculated the average DAI of each sample in both the TCGA and Pri cohort, finding that DAI model failed in predicting the overall survival of IDH wild-type GBMs (Figure 1E-F, Supplementary Figure S3).

**Figure 1.**
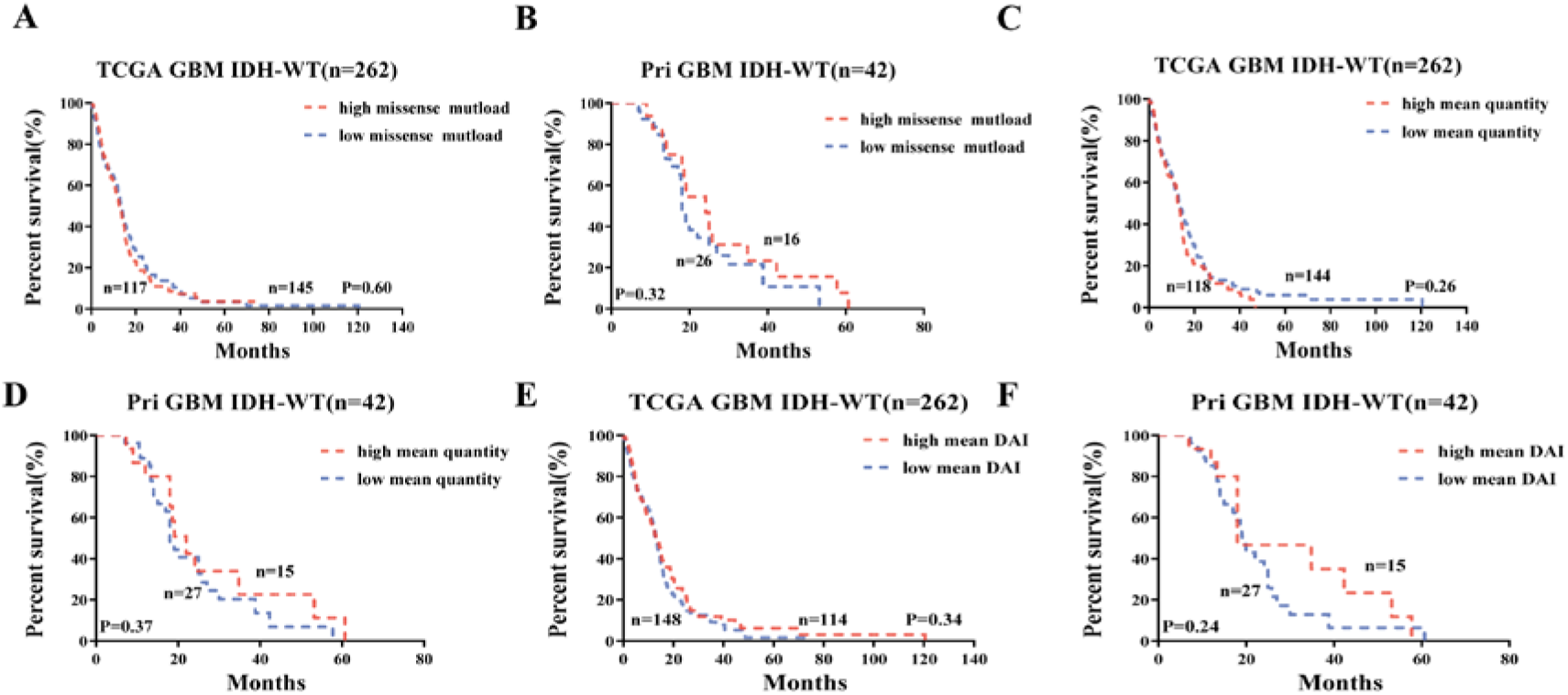
Missense mutational load, absolute number of neoantigens and DAI fail to predict survival of GBM IDH wild-type. A and B, Stratification of survival according to missense mutational load.A, TCGA cohort; B, Pri cohort. red line: high mean mutational load; blue line: low mean mutational load. C and D Stratification of survival according to absolute number of neoanigens; C, TCGA cohort; D, Pri cohort. red line: high mean absolute number of neoantigen; blue line: low mean absolute number of neoantigen. E and F Stratification of survival according to DAI; E, TCGA cohort; F, Pri cohort. red line: high mean DAI; blue line: low mean DAI. n is the number of patients. P-value was determined using log-rank test.

Recent researches emphasize the importance of structural and physical changes relative to self in neoepitope immunogenicity for ovarian cancer[46], and an immunogenic neoantigen must possess structural and physical properties distinct enough to promote efficient recognition by T cells[47]. We first calculated a total of 2928 features for each peptide, mainly including physical-chemical properties, amino acid features, and amino acid descriptors at each absolute position and composed-dipeptide and tripeptide, the site of mutation and the dipeptides and tripeptides related to the mutation site, and complete sequence (Figure 2A). In addition, the Shannon entropy and the AA composition were also calculated. To understand the prognostic effect of each feature, we performed Cox regression to estimate the association between the feature values and overall survival in IDH wild-type GBMs of TCGA. 189 peptide features were predicted to be statistically significantly associated with overall survival (Figure 2B). Among the 189 prognostic features, the most significant positive associations were changes in the absolute site 4 to an aliphatic amino acid (Mutated peptide 4 Aliphatic), ST-scales4 descriptors of site 3 and 4 compose-dipeptide (Mutated peptide 3-4 ST4), and changes in the absolute site 4 to a nonpolar amino acid (Mutated peptide 4 Non.polar). And, the most significant negative associations were theVHSE-scales6 descriptors at the absolute site 4(MT.peptide 4 VHSE6), PP1 descriptors at the absolute site 4(MT.peptide 4 PP1), and polar amino acid at position 4 (MT.peptide 4 polar). To determine the similarities among the 189 prognostic features, we examined their correlation in the TCGA cohort, discovering highly correlated feature modules (Figure 2C). The correlation heatmap of valid features for other glioma subtypes in the TCGA cohort was also calculated, revealing that the correlated feature modules were consistent across different glioma subtypes(Supplementary Figure S4).

**Figure 2.**
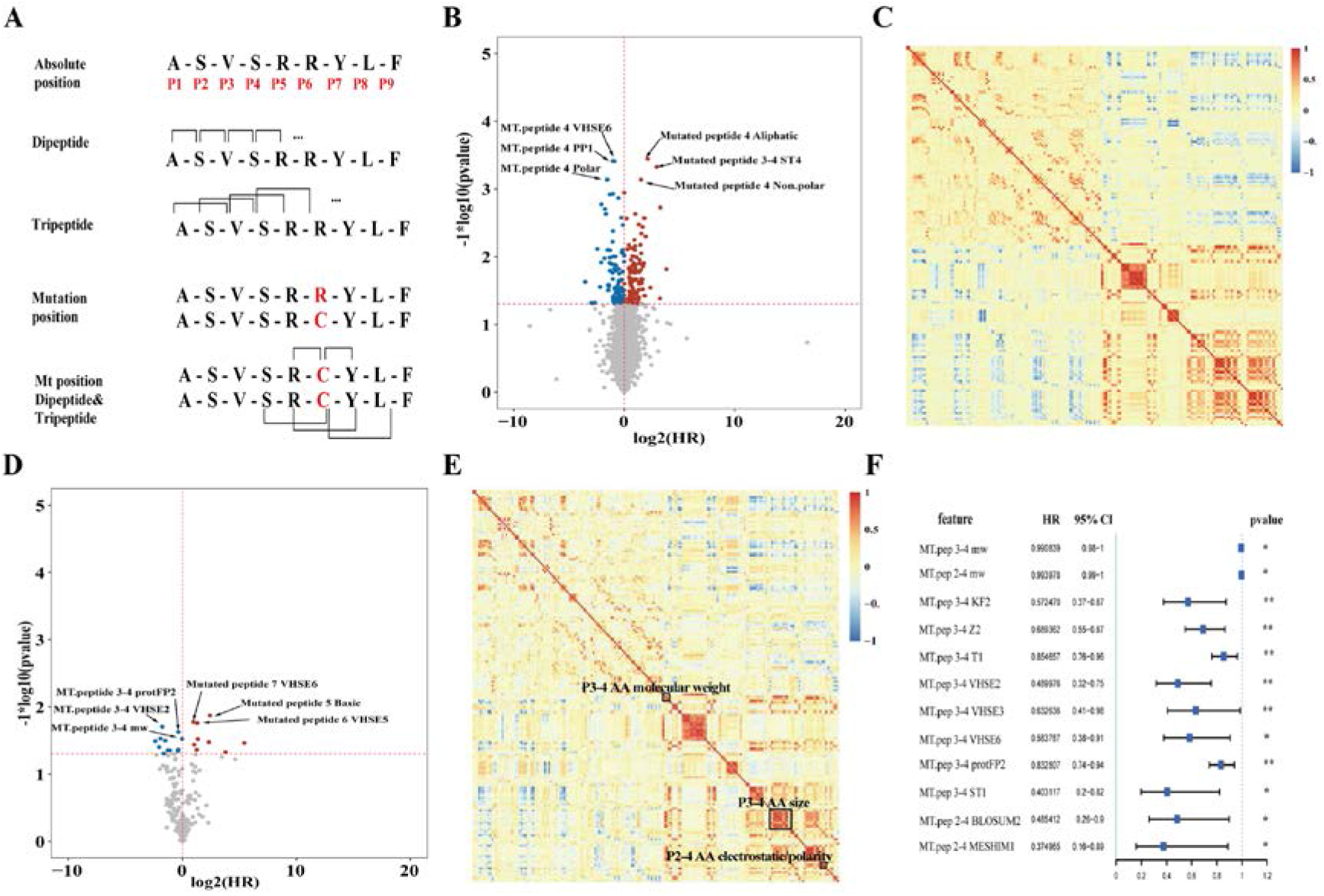
Peptide features associated with the prognosis of patients. A, Summary of the major classes of peptide-intrinsic features identified for each antigen, including amino acid sequence and characteristics at each absolute position, dipeptide, tripeptide, Mutant position, Mutant position dipeptide&tripeptide. Red numbers in position row demonstrate absolute positions of amino acids in peptide. B and D, Volcano plots representing log2(HR) (x-axis) and −log10(pvalue) (y-axis) for each peptide feature as a predictor for prognosis. B, all features in TCGA cohort; D, valid features in Pri cohort. Horizontal dashed line represents the pvalue is equal to 0.05 and the vertical dashed line represents HR is equal to 1. Spot with color represents p-value magnitude lower than 0.05, with the red representing HR above 1 and blue representing HR below 1. C and E, Heat map representing Spearman correlations between each valid feature. C, TCGA cohort; E, Pri cohort. Magnitude of the correlation coefficient represented by color. F, forest plot for 12 peptide features in TCGA cohort.· p-value<0.1;* p-value<0.05;** p-value<0.01. HR value and pvalue were derived by cox regression.

To find out whether these valid features identified from the TCGA database were also of prognostic significance in an independent data of Pri cohort, we conducted Cox regression analysis upon the 189 valid features. A total of 22 features were found to be significantly associated with the overall survival(Figure 2D). The most significant positive associations were VHSE-scales6 at the 7 sites (Mutated peptide 7 VHSE6), changes in the absolute site 5 to a basic amino acid (Mutated peptide 5 basic), VHSE-scales5 at the 6 site(Mutated pep 6 VHSE5). The most significant negative associations were mainly related to the characteristics of the positions 3 and 4 composed-dipeptide, including protFP2 value, VHSE-scales2 value, and molecular weight. Among those features, 12 features were of particular interest as they had shown correlation, and mainly associated with the molecular weight and molecular size/volume of the position 3,4 composed-dipeptide, and molecular electrostatic of the position 2-4 composed-tripeptide(Figure 2E). The 12 features were statistically significant protective factors (hazard ratio < 1) in IDH wild GBMs of the TCGA cohort (Figure 2F) with similar trends observed in Pri cohort (Supplementary Figure S5), demonstrating that neoantigens from long-term survival patients exhibited higher character values.

### Deep learning model using sequence features of neoantigens predicted IDH wild-type GBMs with better survival

Deep learning methods model data by learning high-level representations with multilayer computational models, and rely on algorithms that optimize feature engineering processes to provide the classifier with relevant information that maximizes its performance concerning the final task[22]. Thus, deep learning model is advantageous in learning high-dimensional datasets. Recurrent neural networks(RNN) is a function of all the previous hidden states, but may meet vanishing gradient problem[48, 49]. Long short-term memory(LSTM) can avoid such problem[49], and has the ability to remember all the previous data may help in prediction. To stratify IDH wild-type GBMs using the 189 features, we constructed a deep learning model including three hidden layers (two LSTM layers and one fully connected layer) with 128, 32, 8 nodes, respectively(Figure 3A). We chose the Sigmoid function as neuron activation function for fully connected layer, MSE as the loss function and Adam as the iterative optimizer. Considering the relationship between the number of iterations and the loss function of the model, the final number of iterations was selected as 1000(Supplementary Figure S6). Initially, the connection weights and biases of each layer were randomly generated. The samples in the TCGA cohort were used as training data, while the samples in Pri cohort were used as external testing data.

**Figure 3.**
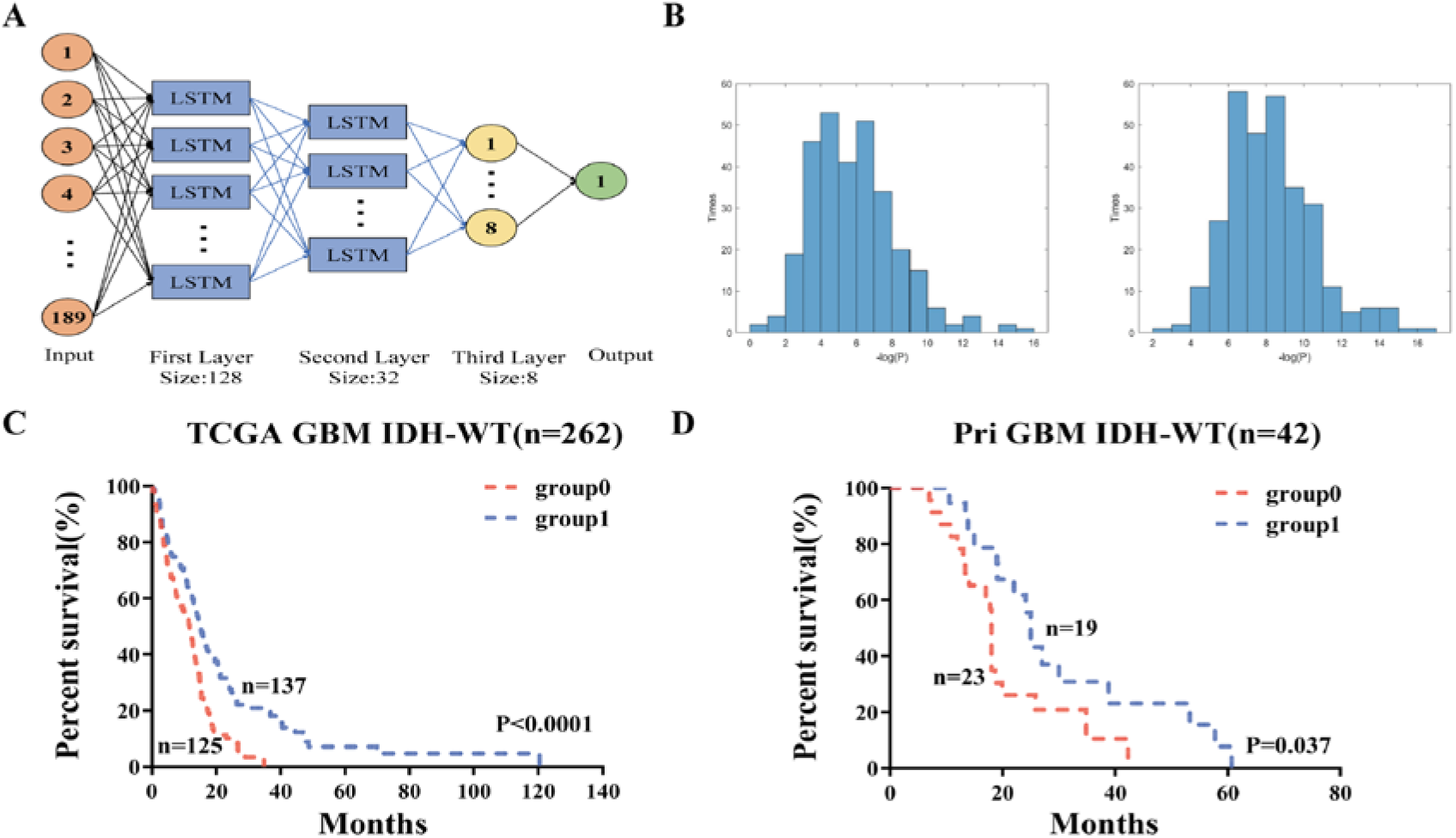
Deep learning model predict better survival in GBM. A, deep learning model diagram. B, Left, pvalue distribution representing −log(pvalue)(x-axis) and times(y-axis) for 300 times in cross validation. The data used for verification was 40% and 60% of TCGA samples, respectively. Right, survival of 60% TCGA patients stratified according to the final model. C, Survival of TCGA cohort patients stratified according to the final model. D, Survival of Pri cohort patients stratified according to the final model. red line, the deep learning model prediction label is 0; blue line, model prediction label is 1. P-value was performed using log-rank test. n is the number of patients.

To validate the reliability of the deep learning model, we performed 300 random trials with each splitting the samples into training set and internal testing set at the ratio of six vs four. In each trial, the parameters learned in the training set were applied in the internal testing set. In 275 out of 300 trials, IDH wild-type GBMs in TCGA were successfully separated into two prognostic subgroups with the overall survival significantly different (P < 0.05) (Figure 3B left), demonstrating the high stability of our model. The optimal parameter settings were determined from the 275 successful trials and applied to randomly selected 60% of IDH wild-type GBMs in TCGA. In 299 of 300 randomly selected 60% of IDH wild-type GBMs in TCGA, the deep learning model with the optimal parameter successfully separated patients into two significantly prognostic subgroups (Figure 3B right), demonstrating that the model we built can predict IDH wild-type GBMs with better survival in TCGA stably and reliably. With the use of the optimal model, all IDH wild-type GBMs in TCGA were separated into two significantly prognostic subgroups (P<0.0001, Figure 3C). We applied the optimal model in an independent data (Pri GBM cohort), finding that patients in Pri GBM cohort were successfully separated into two subgroups with statistically significant overall survival (*P* = 0.037) (Figure 3D). We also applied the optimal model to subgroups in the TCGA pan-glioma cohort, revealing that in GBM, IDH wildtype, Classical, Classical-like, Mesenchymal-like subtypes, patients grouped by the model demonstrated a significant difference in overall survival (*P* < 0.05) (Supplementary Figure S7).

### The prognostic characteristics of 12 protective sequence features

To characterize the 12 protective sequence features in the molecular weight, molecular size of dipeptide, and molecular electrostatic potential of tripeptide, we compared their distributions in the short- and long-term survival IDH wild-type GBMs. In comparison with the short-term survival IDH wild-type GBMs, the long-term survival patients exhibited statistically significantly higher molecular weight of dipeptide at the site 3 and 4 (*P*<0.05; Figure 4A; Supplementary Figure S8a), molecular size-related features (Kidera Factors 2, Z-scale 2, T-scale 1, protFP2, VHSE-sclae 2, VHSE-sclae 3, VHSE-sclae 6, ST-scale 1) (*P*<0.05; Figure 4B; Supplementary Figure S8b) and the electrostatic potential related features (the BLOSUM2 value and the MESHIM1 value) (*P*<0.05; Figure 4C; Supplementary Figure S8c) in both TCGA and Pri-cohort.

**Figure 4.**
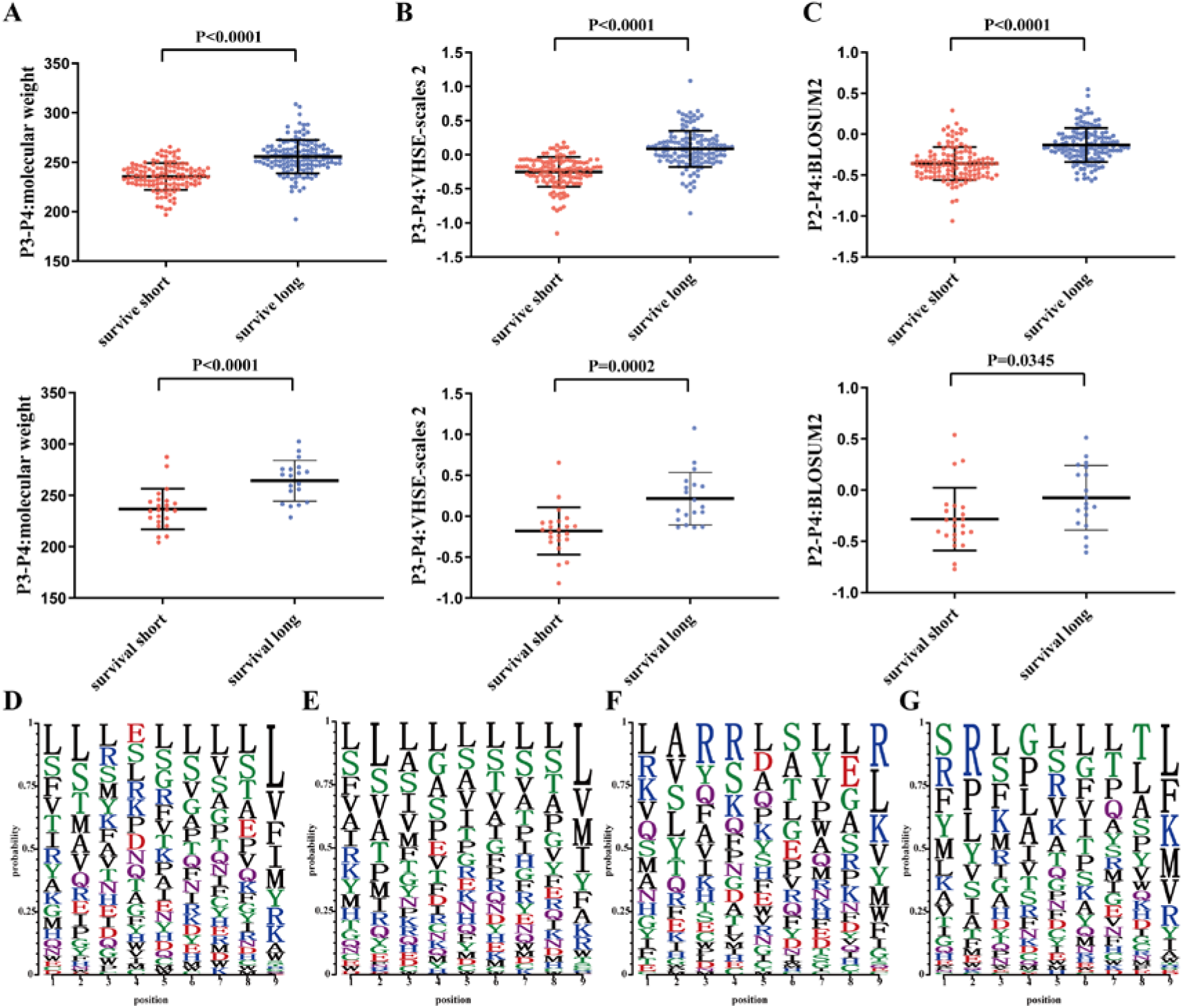
Annotation of features associated with prognosis. A-C, Comparison of the similarity between valid features value in long-term survival and short-term survival groups of IDH wild-type GBM in two cohorts. A, molecular weight of dipeptide composed with the site 3 and 4. B, VHSE-scales2 of dipeptide composed with the site 3 and 4. C, BLOSUM2 of tripeptide composed with the site 2 and 4. The upper and the lower row contain the TCGA cohort and Pri cohort, respectively. P value was calculated by unpaired T test. D-G, Comparison of the amino acid occurrence frequency for each absolute position between the two groups predicted by deep learning model. D, long-term survival patients in TCGA cohort; E, short-term survival patients in TCGA cohort; F, long-term survival patients in Pri cohort; G, short-term survival patients in Pri cohort. the higher the occurrence, the taller the letter.

Univariate and multi-variate Cox regression analysis demonstrated that two of twelve sequence features (VHSE2 value and protFP2 value) were associated with the overall survival in both IDH wild-type GBM of TCGA cohort and Pri cohort (Supplementary Table S1-S4). Kaplan Meier analysis demonstrated that there was statistically significantly different overall survival between the low-value (below mean) and high-value (above mean) groups of IDH wild-type GBMs stratified by the two features. The patients with high value (above the mean values) had a significantly longer overall survival (for protFP2 value: *P* = 0.002 in TCGA cohort and P=0.03 in Pri cohort; for VHSE2 value: *P* = 0.018 in TCGA cohort and P=0.11 in Pri cohort) (Supplementary Figure S9a-b). Furthermore, the two feature-based stratification of the IDH wild-type GBMs were found independent of age and mutational load. In addition, the two features exhibited strong correlations (R = 0.87, P < 2.2e-16 for TCGA; R= 0.91, P < 2.2e-16 for Pri Cohort) (Supplementary Figure S9c).

The distributions of amino acid residue for 9-mers between long- and short-term survival groups of IDH wild-type GBMs were examined, revealing that the ratios of amino acid residues at positions 3 and 4 were significantly different (Figure 4D-G). At the site 3, the patients with neoantigens containing a lower frequency of L and S amino acid and a higher frequency of R amino acid survived longer than those with the opposite frequencies in both the TCGA cohort and Pri cohort. The enrichment of residues R and S at site 4 of neoantigens were evident in the long-term survival of IDH wild-type GBMs. The ratios of L and G at site 4 of neoantigens increased in the short-term survival patients.

### Tumor Purity and functional annotation of gene expression in GBM

Recent researches showed that the tumor purity estimated from the DNA, RNA and methylation-based methods had high concordance among most cancer types[31][48]. To compare the difference of tumor purity between long-term and short-term survival groups of IDH wild-type GBMs, we calculated the tumor purity values for each patient in both TCGA and Pri cohorts. No significant difference in tumor purity was observed between long- and short-term survival of IDH wild-type GBMs (Figure 5A, Figure 5B). We also found that there was no significant difference in regards to immune scores and stromal scores between long- and short-term survival groups (Supplementary Figure S10a-b), suggesting that immune cell infiltrations were prognostic in IDH wild-type GBMs. No correlations were discovered between purity levels and mutational burden (Supplementary Figure S10C).

**Figure 5.**
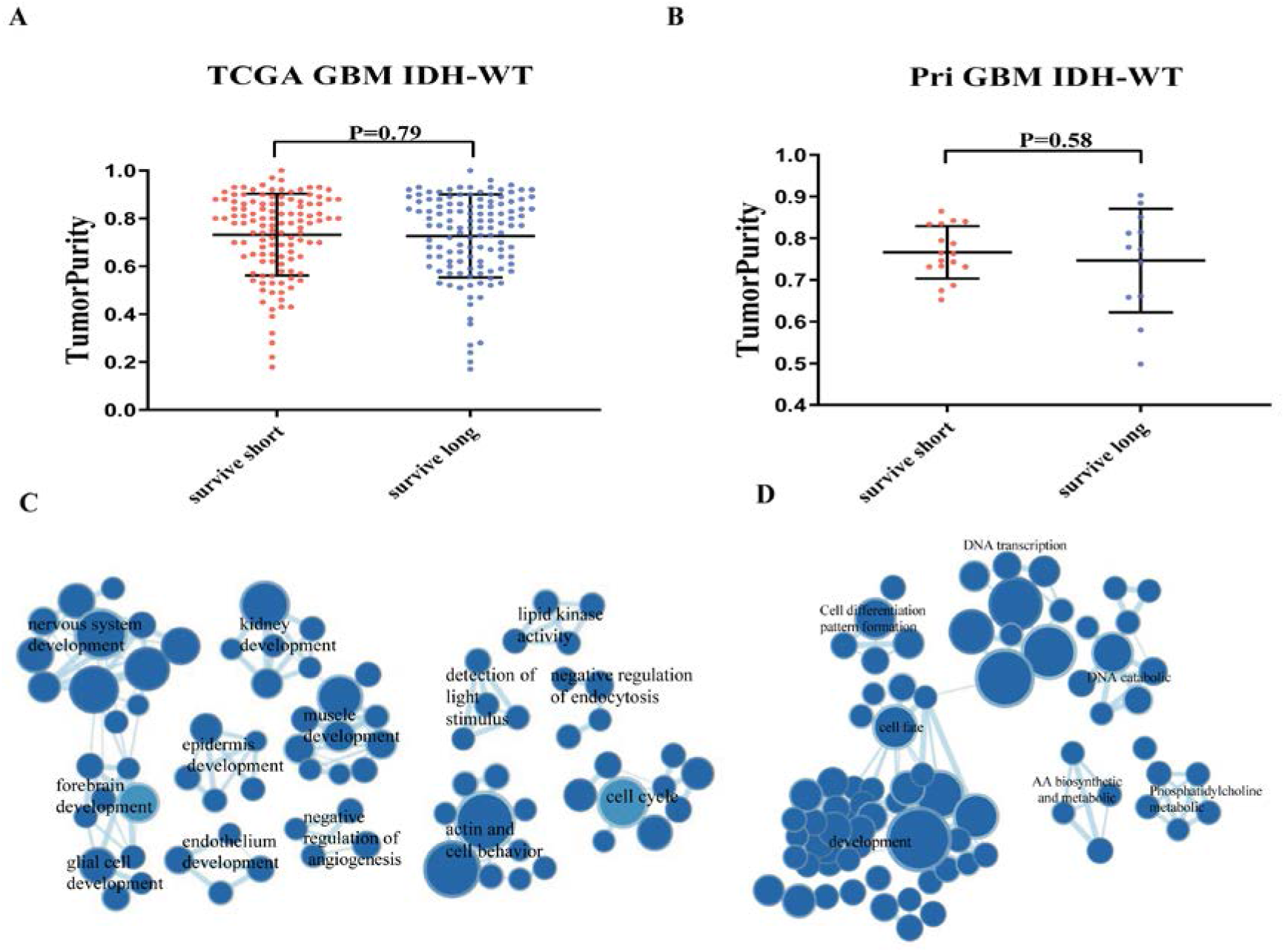
Tumor Purity and functional annotations of gene expression in GBM. A and B, Comparison of the similarity between tumor purity in long-term survival and short-term survival groups of IDH wildtype GBM in two cohorts. A, TCGA cohort. B. Pri cohort. P value was calculated by T test. C and D, Enrichment map network of statistically significant GO categories in the patient cohort with p-value<0.05. C, TCGA cohort. D. Pri cohort. Nodes represent GO terms and lines represent their connectivity. Node size is proportional to the number of genes in the GO category and line thickness indicates the fraction of genes shared between groups.

To understand the mechanisms in transcriptomic architecture, Gene Set Enrichment Analysis (GSEA) was conducted between long- and short-term survival group of IDH wild-type GBMs in both TCGA and Pri cohorts, respectively. Enrichment map analysis of deregulated GO terms in TCGA data demonstrated that GO terms related to nervous development, glial cell development, epidermis development, cell cycle, muscle tissue development, and kidney development were highly enriched in long-term survival patients (Figure 5C, Supplementary Table S5-S6). For Pri cohort, the most significant biological processes enriched in longer-survived patients were development associated GO terms such as epidermis development, cell cycle, which were also identified in IDH wild-type GBMs in the TCGA cohort (Figure 5D).

## Discussion

In this paper, we provided a prognostic prediction method based on neoantigen peptide-intrinsic features. Although several survival prediction models have been reported based on the expression of a few genes [50–52]or medical images [53, 54] with successfully predicting survival in patients with glioblastoma, those features used in the methods are not related to immune response. Therefore, the above predictions could not predict immunoreaction. Since characters extracted from noeantigens are associated with tumor-specific T-cell responses and anti-tumor immune responses, the method we provided in this article can help predict the prognosis of IDH wild-type GBM patients who will likely benefit from neoantigen based personalized immunetherapy.

An LSTM model was proposed to improve predictive performance by integrating neoantigen features into deep learning model, and applied to the survival prediction of IDH wild-type GBM patients. Results showed the model obtained better predictive performance in two data cohort, and performed excellent prediction in some higher-grade glioma subtypes, including Classical, Classical-like, Glioblastoma, IDHwildtype, Mesenchymal-like. Additionally, we also identified two correlated neoantigen features, which stratified patients into a high- and low-value group with significant survival difference with independent of other clinical and pathological features.

The analysis of valid features demonstrated their internal connections. Among these, 12 features associated with better survival status were analyzed, mainly including amino acid molecular weight, molecular size/volume, and electrostatic potential/polarity. Additionally, the 12 features performed close relation with the amino acid properties at the absolute positions 3 and 4 of the mutant peptide, which is also confirmed using the amino acid distributions between the different survival status groups. The features at the site 3 and 4 of the neoantigen may therefore have potential effect on the survival of GBM patients and immunotherapy response, and they are worthy of further investigation.

The LSTM network originated from RNN and developed by exploding and vanishing gradients while training traditional RNN networks. It could learn potentially effective information from valid neoantigen features defined with remarkable flexibility and adaptability and help predict prognosis. The model informed a cohort of patients who have better clinical prognosis result, which generally exhibits biological process such as development and cell cycle. It is well known that GBM typically lacks a significant number of T lymphocyte infiltrates[55]. Lymphocyte depletion and immunosuppressive microenvironment are distinctive features of malignant gliomas[56]. Thus, we also did tumor purity analysis. On the opposite to feature correlations analysis, tumor purity analysis showed no significant relation with different survival status group, indicating that patients survival status are independent of broad immune cell infiltration.

Comparing to traditional cytological identification of cancer, the LSTM model is superior because it won’t be influenced by diagnostic instruments, and machine learning is more effective in the analysis of complex data. Our models are still facing some limitations. For instance, we mainly focused on protein primary structure in this study, but did not place enough emphasis on secondary and tertiary protein structure. Thus, more features might be integrated into the model to promote prediction accuracy. Those issues shall be resolved in the future. The deep learning method could be used to augment the prognostic evaluation and improve decision-making in glioma. To predict the patients’ outcome, more studies related to generalizability test are still in need.

## Methods

### Feature calculation for neoantigens

For the purpose of extracting features from neoantigens, the samples with detected mutant peptides remained in the downstream analysis, including 262 samples in the TCGA cohort and 42 samples in Pri cohort. A total of 2928 features were extracted from 2263 neoantigens (2081for TCGA cohort; 182 for Pri cohort) in the downstream analysis. Specifically, features used in the calculation were derived using the R package “Peptides”(v2.4.2) including 66 amino acid descriptors and physical-chemical properties (aliphatic, auto-correlation, auto-covariance, Boman index, theoretical net charge, cross-covariance, hydrophobic moment, hydrophobicity, instability, molecular weight). Additionally, the “aaComp” command was also used to describe amino acid features including Tiny, Small, Aliphatic, Aromatic, Non-polar, Polar, Charged, Basic, Acidic. Variables were derived by the presence (1) or absence (0) of each feature. Characteristic variables were performed in four conditions respectively, including the complete sequence, the site of mutation along with each antigen and the dipeptides/tripeptides related to the mutation site, each absolute position along each antigen and related dipeptide/tripeptide composition, and the difference of each feature in the mutated versus reference antigen.

The features, described overall content of a protein, for example, amino acid composition, were significant. Variables demonstrating presence(1) or absence(0) of each amino acid type following, including the first or last 3 amino acid residues or middle residues of each antigen, the first or last amino acid residues of each antigen, the first or last 2 amino acid residues or middle residues of each antigen.

To measure the complexity at the protein and residue level, we computed Shannon entropy of a protein and entropy of each type of residues using the following equations:

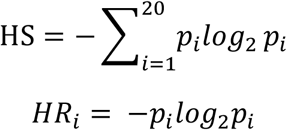

where HS is Shannon entropy of a protein sequence and *HR_i_* is the entropy of a residue type i. p*_i_* is the probability of the existence of a given amino acid in the sequence. We calculated the Shannon entropy of the mutant peptides and the difference of Shannon entropy in the mutant antigen versus reference antigen. Cancer is characterized by the accumulation of mutations, so the analysis of mutant positions is valid. Therefore, the Shannon entropy of the dipeptides/tripeptides related to the mutation site and the entropy difference of mutations process were performed. The entropy of a residue type was also calculated for the mutant peptides and reference peptides.

### Prognostic feature selection

The features were calculated for all detected 9-mer mutated peptides and wild peptides. There were multiple types of mutations in each patient, resulting in a large number of mutant peptides in total. Thus, each feature value calculated in all the peptides detected in a patient was averaged as the final value. Univariate Cox regression analysis was performed here to predict the impact of each feature on prognosis. The threshold of P-value was set as 0.05, which means all the features with P-value lower than or equal to 0.05 were deemed as statistically significant (termed as valid features). A correlation matrix of the valid features was conducted, and visualized through heatmaps using the package ‘pheatmap’ in R language.

### Hierarchical k-means clustering

Hierarchical k-means clustering was applied upon the Z-Score-transformed valid features to stratify patients into two clusters. Hierarchical k-means was performed using the “hkmeans” command of the R package ‘factoextra’ (version 1.0.7). The overall survival differences between two clusters of patients were compared through Kaplan-Meier survival analysis.

### Deep-learning model construction

The grouping results derived from hierarchical k-means clustering were used as labels, marking 0 and 1. The valid features in the TCGA cohort were used as training data to train the deep learning model. The input data were Z-Score-transformed valid features, in order to avoid gradient disappearance problem. The LSTM deep learning model was built with three hidden layers, including two LSTM layers and one fully connected layer, each layer containing 128, 32, and 8 nodes, respectively. Sigmoid function was chosen as neuron activation function for fully connected layer, MSE as the loss function and Adam as the iterative optimizer. The maximum number of iterations was set as 1000. The initial connection weights and biases of each layer were randomly generated, and end up reaching stable parameters through training iterations.

### Leave one out cross validation (LOOCV)

After determining the framework of the model, cross validation was a necessary step. Specifically, the training data was separated into two sections randomly with proportion of training and testing sets as 6 to 4. The training set was used to train the model to determine the unknown parameters, while the test set was used to validate the effect of the predicted parameters. To obtain the optimal model, the above process was carried out 300 times. Kaplan-Meier survival analysis was operated each time to see if the model can divide the samples into two groups with a statistically significant survival difference. Only groups with P-value lower than or equal to the threshold of 0.05 were regarded as statistically significant. Among 300 times trial, the more significant stratifications, the more stable our model is.

### Independent validation

A model with fixed parameters corresponding to the lowest P-value was selected as the optimal model. To test the performance of the optimal model, Pri cohort was used as an external test data. The optimal model divided patients in the Pri cohort into long- and short-term survival clusters. Kaplan Meier analysis was conducted between the long- and short-term survival clusters in Pri cohort to test the predictive performance of the optimal model for IDH wild-type GBMs. Besides, other glioma subtypes from TCGA data were also tested by the model, including Astrocytoma, Classical-like, Classical, Codel, Glioblastoma, G-CIMP-high, IDH-MT-codel, IDH-MT-noncodel, IDH-MT, IDH-WT, Mesenchymal-like, Mesenchymal, Neural, Oligodendroglioma, Proneural and OligoAstrocytoma.

### Tumor purity estimation

Tumor purities were estimated by ESTIMATE[57] from gene expression profiles. There were a total of 242 and 29 IDH wild-type GBMs in the TCGA cohort and Pri cohort with gene expression profiles available, respectively. The purity score was performed for each sample, using the R package ‘estimate’(version 1.6.7). Meanwhile, the immune score and the stromal score were also estimated.

### GO enrichment analysis

To identify differential genes expression between different groups, GO enrichment analysis was conducted using Gene Set Enrichment Analysis (GSEA 4.0.3)[58], with 17814 and 23491 genes available in the TCGA cohort and Pri cohort, respectively. The GO terms were collected from the Molecular Signatures Database (c5.all.v6.2.symbols.gmt), including cellular component, molecular function, and biological process. Number of Permutations was set to 1000 and gene set size filters were 15-500. Gene sets with FDR <0.05 were considered as differentially expressed, and visualized using Cytoscape[59]. The grouping results of GO terms were shown in Supplementary Table S5-S6.

### Statistical Analysis

All statistical analyses were performed using R software, version 4.0.0. Continuous variables between groups were compared by the unpaired T test. Correlations between continuous variables were evaluated by Pearson correlation analyses. For all statistical analyses, the P value of 0.05 was taken as the significant threshold in all tests. Kaplan-Meier survival analysis curves were compared using a log-rank test and Multivariate survival analysis was performed by Cox regression model using R package “survminer” and “survival”. All analyses were conducted in R language and Python language.

## Availability of Source Code and Requirements

Project name: Neoantigen-intrinsic feature based Deep Learning Model

Project home page: https://github.com/zhangjbig/neoDL

Operating system: Platform independent

Programming language: Python, R

Other requirements: Python 3.8 or higher, R 3.3 or higher

License: GPL-2

## Availability of Supporting Data and Materials

All data used for this article are available at the following websites or accession numbers: (i) Mutations and clinical information in the TCGA cohort[36]. (ii) Gene expression microarray data with Agilent chip (G4502A) at level 3: TCGA Data portal. (iii) Mutations, RNAseq gene expression data, and clinical information in the Pri cohort[37]. (iv) The intrinsic features of neoantigens for each sample in both TCGA cohort and Pri cohort are available at github (https://github.com/zhangjbig/neoDL).

## Additional Files

Supplementary Figure S1. Survival of glioma patients stratified according to missense mutational load.

Supplementary Figure S2. Survival of glioma patients stratified according to absolute number of neoantigens.

Supplementary Figure S3. Survival of glioma patients stratified according to differential agretopicity index(DAI).

Supplementary Figure S4. Heatmap representing Spearman correlation between each valid feature.

Supplementary Figure S5. Forest plot for 12 peptide features in Pri cohort.

Supplementary Figure S6. Relationship between the number of iterations and loss/accuracy.

Supplementary Figure S7. Survival of TCGA glioma patients stratified by deep learning model.

Supplementary Figure S8. Comparison of the similarity of valid feature values between long-term survival and short-term survival groups of IDH wild-type GBM in two cohorts.

Supplementary Figure S9. Survival of glioma patients stratified according to 2 feature values, and analysis of the correlations between these features in two cohorts.

Supplementary Figure S10. Comparison of the similarity of immune score and stromal score between two groups and the correlation analysis between purity and mutation load.

Supplementary Table S1. Multivariate Cox regression analysis including position 3-4 composed-dipeptide VHSE-scale 2 value, mutation load and age for TCGA IDH wild-type GBM (n=262).

Supplementary Table S2. Multivariate Cox regression analysis including position 3-4 composed-dipeptide VHSE-scale 2 value, mutation load and age for Pri IDH wild-type GBM (n=42).

Supplementary Table S3. Multivariate Cox regression analysis including position 3-4 composed-dipeptide protFP 2 value, mutation load and age for TCGA IDH wild-type GBM (n=262).

Supplementary Table S4. Multivariate Cox regression analysis including position 3-4 composed-dipeptide protFP 2 value, mutation load and age for Pri IDH wild-type GBM (n=42).

Supplementary Table S5. Functional annotation for the lists of genes differentially expressed analyzed by GSEA in TCGA cohort.

Supplementary Table S6. Functional annotation for the lists of genes differentially expressed analyzed by GSEA in Pri cohort.

## Abbreviations

GBM: glioblastoma
LSTM: Long short-term memory
RNN: Recurrent neural networks
GSEA: gene set enrichment analysis.

## Competing interests

The authors declare no competing interests.

## Funding

This work was supported by grants from the National Natural Science Foundation of China (NSFC No.11421202, and 11827803 to YBF, No.81672479 to W.Z), National Natural Science Foundation of China (NSFC)/Research Grants Council (RGC) Joint Research Scheme (81761168038)(W.Z.), Beijing Municipal Administration of Hospitals’ Mission Plan (SML20180501)(W.Z.), the Youth Thousand Scholar Program of China (J.Z.) and Program for High-Level Overseas Talents, Beihang University (J.Z.).

## Author Contributions

Conceptualization: J.Z., W.Z., Y.B.F.; Methodology: T.S., Y.F.H., W.D.L., G.L., J.Z., W.Z., Y.B.F.; Data Curation: G.L., L.L., W.D.L., L.W., Z.X.X., X.H.H., H.W., Y.L., Y.F.C., H.Y.W.,and J.L.; Writing-Review and Editing: T.S., Y.F.H., Z.X.X., W.Z., Y.B.F., J.Z.; Supervision: W.Z.,Y.B.F.,J.Z.; Funding Acquisition: W.Z., Y.B.F.,J.Z.

## Notes

### Competing Interest Statement

The authors have declared no competing interest.

