## Supplementary Materials for "neoDL: A novel neoantigen intrinsic feature-based deep learning model identifies IDH wild-type glioblastomas with the longest survival"

**Supplementary Materials Legends**

**Supplementary Figure S1.** Survival of glioma patients stratified according to missense mutational load.

**Supplementary Figure S2.** Survival of glioma patients stratified according to absolute number of neoantigens.

**Supplementary Figure S3.** Survival of glioma patients stratified according to differential agretopicity index (DAI).

**Supplementary Figure S4.** Heat map representing Spearman correlation between each valid feature.

**Supplementary Figure S5.** Forest plot for 12 peptide features in Pri cohort.

**Supplementary Figure S6.** Relationship between the number of iterations and loss/accuracy.

**Supplementary Figure S7.** Survival of TCGA glioma patients stratified by deep learning model.

**Supplementary Figure S8.** Comparison of the similarity of valid feature values between long-term survival and short-term survival groups of IDH wild-type GBM in two cohorts.

**Supplementary Figure S9.** Survival of glioma patients stratified according to 2 feature values, and analysis of the correlations between these features in two cohorts.

**Supplementary Figure S10.** Comparison of the similarity of immune score and stromal score between two groups and the correlation analysis between purity and mutation load.

**Supplementary Table S1.** Multivariate Cox regression analysis including position 3-4 composed-dipeptide VHSE-scale 2 value, mutation load and age for TCGA IDH wild type GBM (n=262).

**Supplementary Table S2.** Multivariate Cox regression analysis including position 3-4 composed-dipeptide VHSE-scale 2 value, mutation load and age for Pri IDH wild type GBM (n=42).

**Supplementary Table S3.** Multivariate Cox regression analysis including position 3-4 composed-dipeptide protFP 2 value, mutation load and age for TCGA IDH wild type GBM (n=262).

**Supplementary Table S4.** Multivariate Cox regression analysis including position 3-4 composed-dipeptide protFP 2 value, mutation load and age for Pri IDH wild type GBM (n=42).

**Supplementary Table S5.** Functional annotation for the lists of genes differentially expressed analyzed by GSEA in TCGA cohort.

**Supplementary Table S6.** Functional annotation for the lists of genes differentially expressed analyzed by GSEA in Pri cohort.

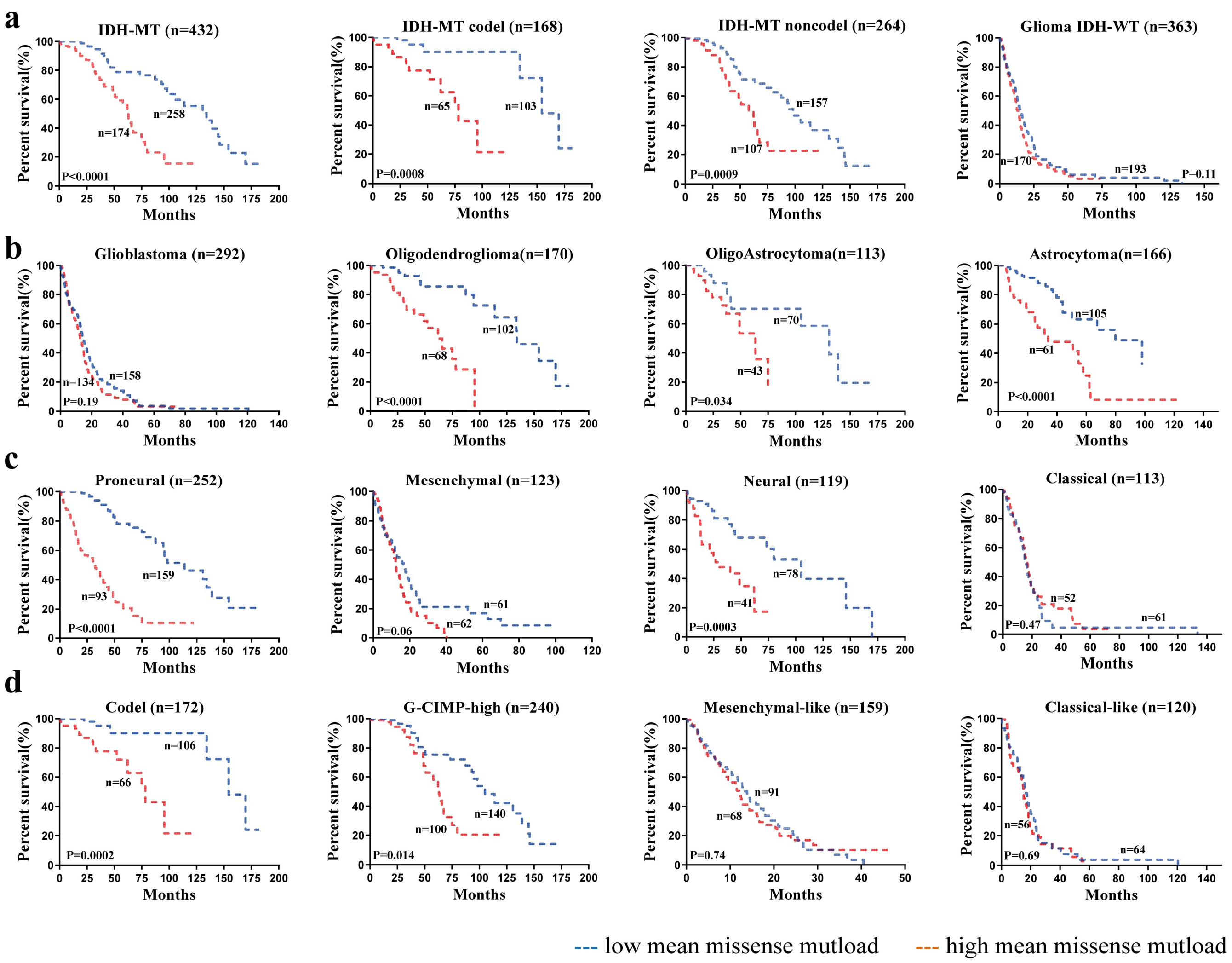


**Supplementary Figure S1.** Survival of glioma patients stratified according to missense mutational load. **a**, Glioma IDH status sub-groups; **b**, Glioma histology sub-groups; **c**, Glioma transcriptomic sub-groups; **d**,Glioma DNA methylation sub-groups. red line, high mean mutational load ; blue line, low mean mutational load; n, number of patients; p-value was determined using the log-rank test.

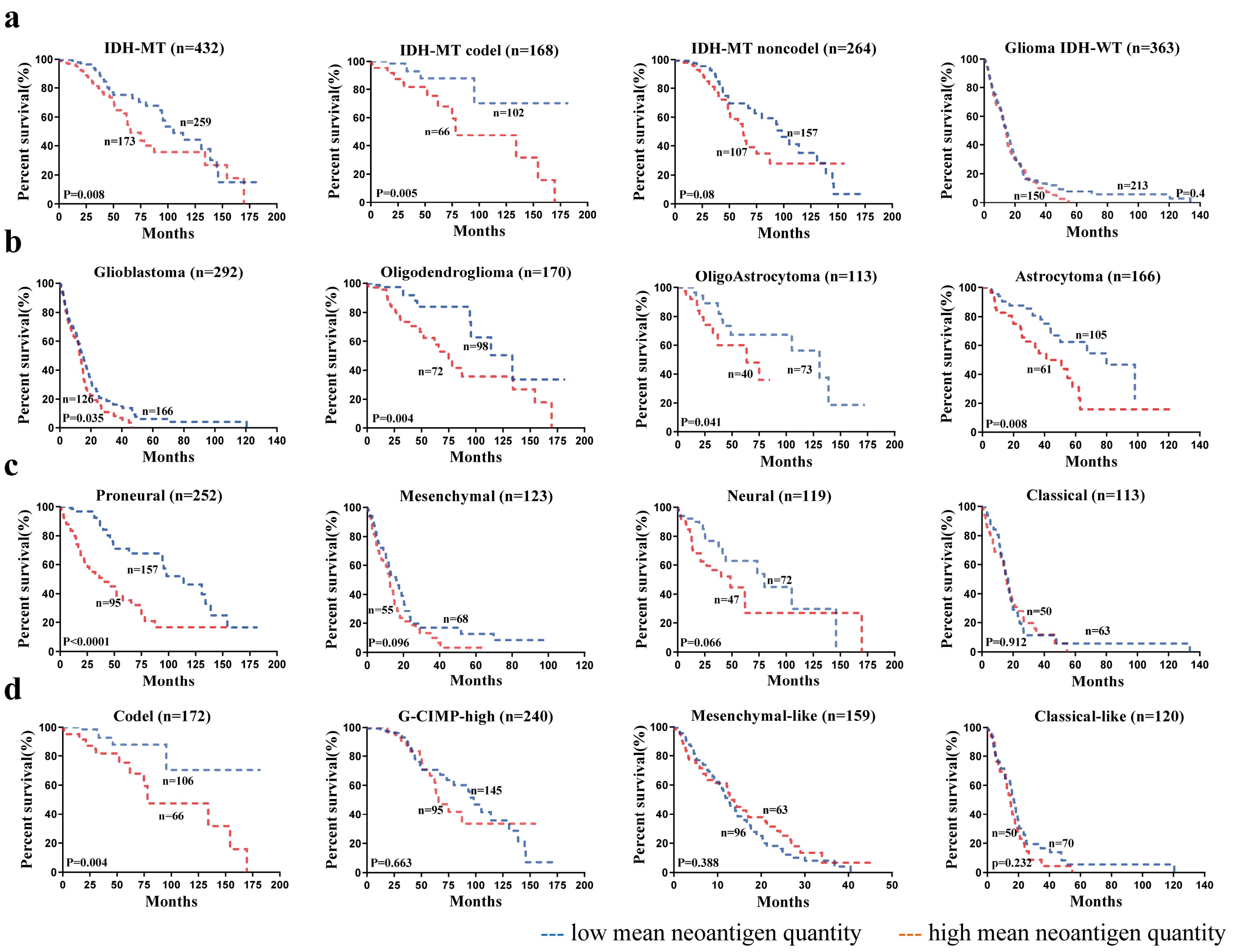


**Supplementary Figure S2.** Survival of glioma patients stratified according to absolute number of neoantigens. **a**, Glioma IDH status sub-groups; **b**, Glioma histology sub-groups; **c**, Glioma transcriptomic sub-groups; **d**,Glioma DNA methylation sub-groups. red line, high mean mutational load ; blue line, low mean mutational load; n, number of patients; p-value was determined using the log-rank test.

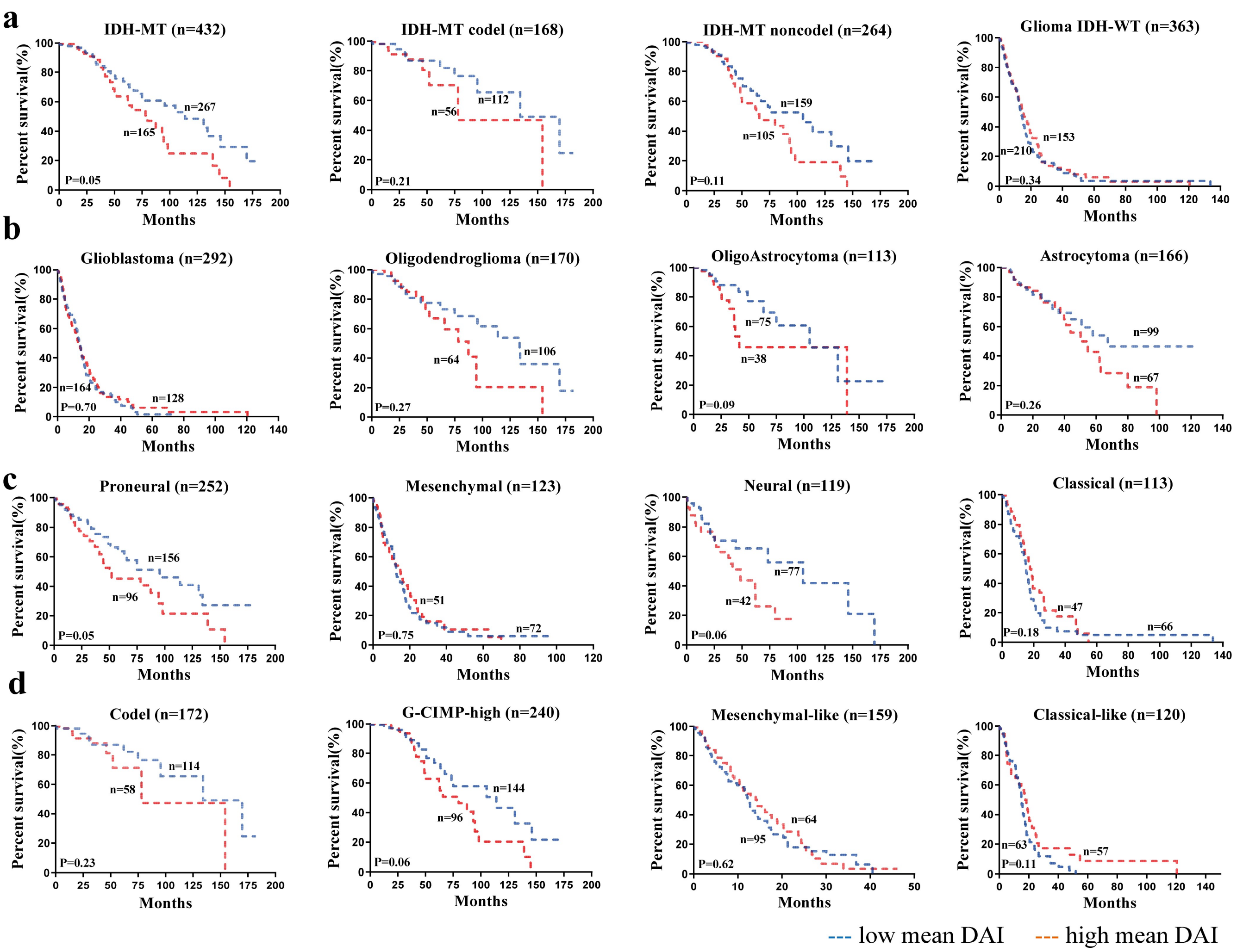


**Supplementary Figure S3**. Survival of glioma patients stratified according to differential agretopicity index (DAI). **a**, Glioma IDH status sub-groups; **b**, Glioma histology sub-groups; **c**, Glioma transcriptomic sub-groups; **d**,Glioma DNA methylation sub-groups. red line, high mean DAI; blue line, low mean DAI; n, number of patients; p-value was determined using the log-rank test.

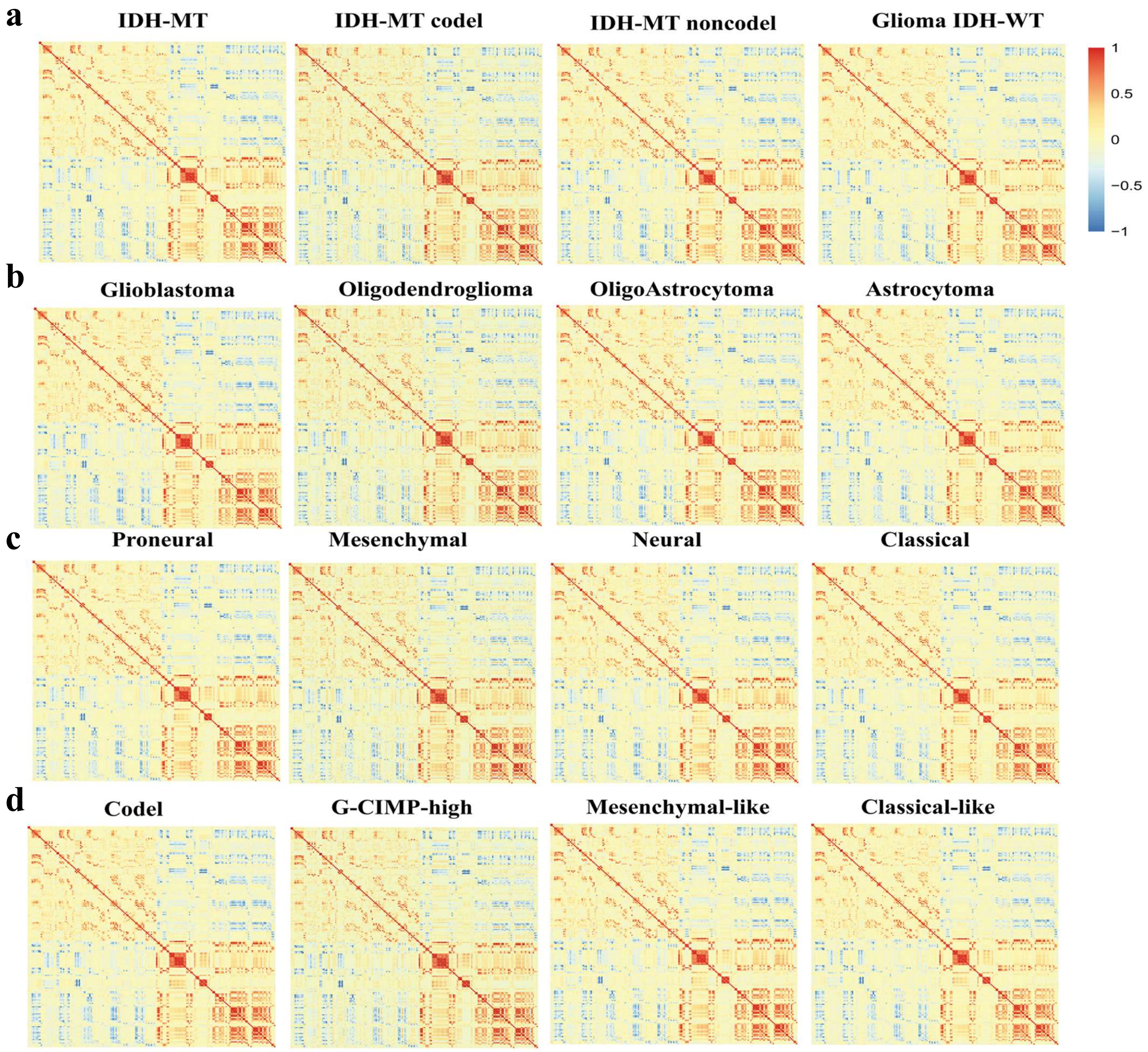


**Supplementary Figure S4**. Heat map representing Spearman correlations between each valid feature. Magnitude of the correlation coefficient represented by color. **a**, Glioma IDH status sub-groups; **b**, Glioma histology sub-groups; **c**, Glioma transcriptomic sub-groups; **d**,Glioma DNA methylation sub-groups.

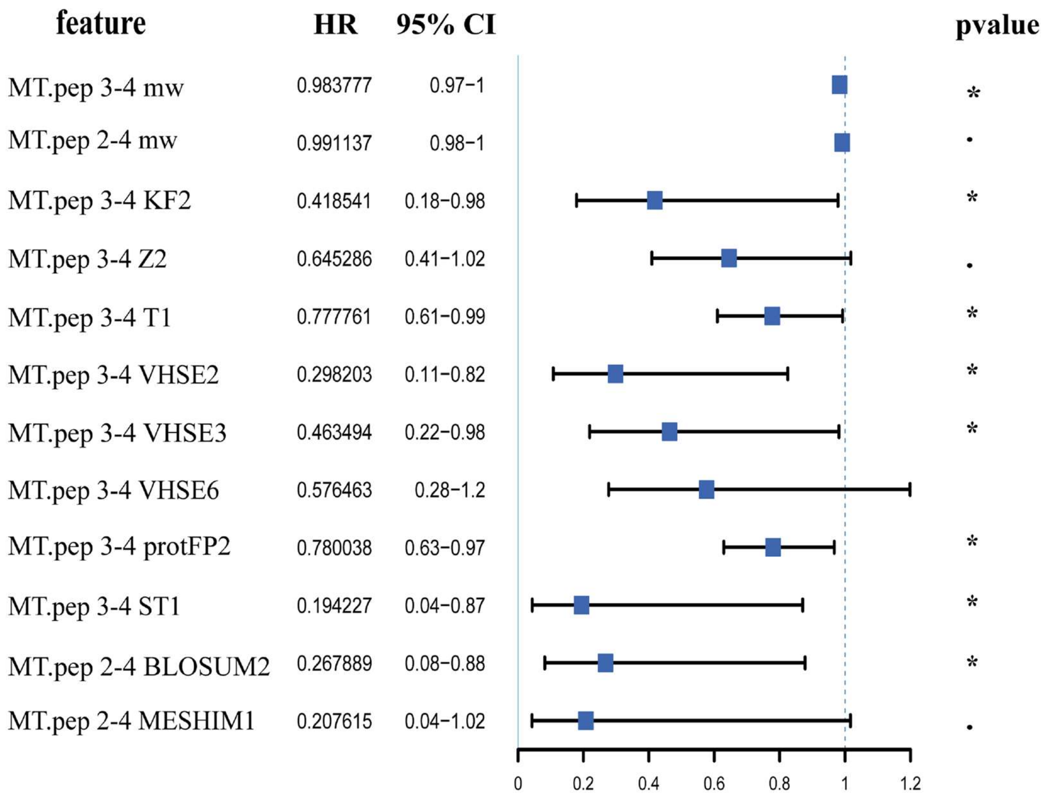


**Supplementary Figure S5.** forest plot for 12 peptide features in Pri cohort. · pvalue<0.1;* pvalue<0.05;** pvalue<0.01. HR value and pvalue derived by cox regression.

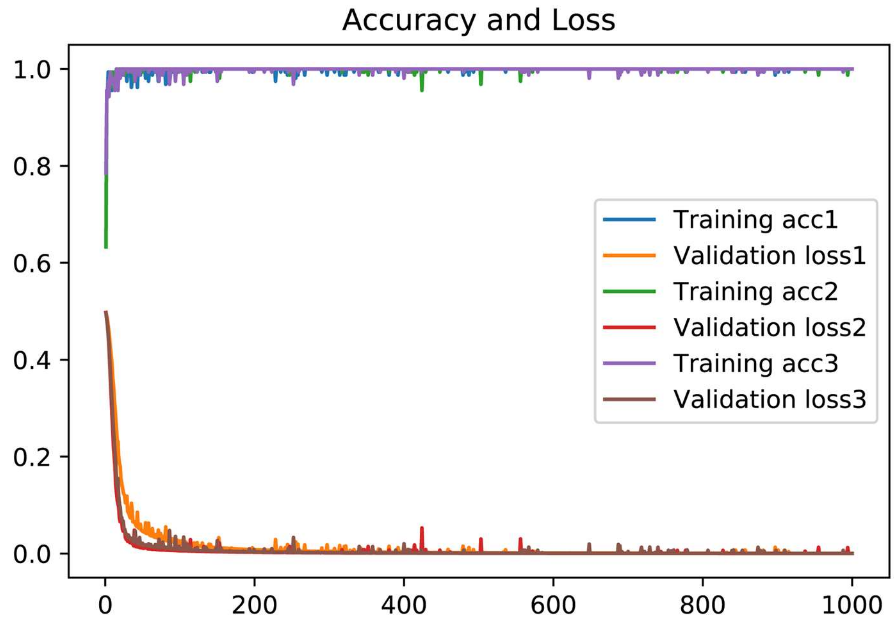


**Supplementary Figure S6**. Relationship between the number of iterations and loss/accuracy. The number of iterations is set to 1000, and the loss/accuracy change curve in the three training processes is randomly selected.

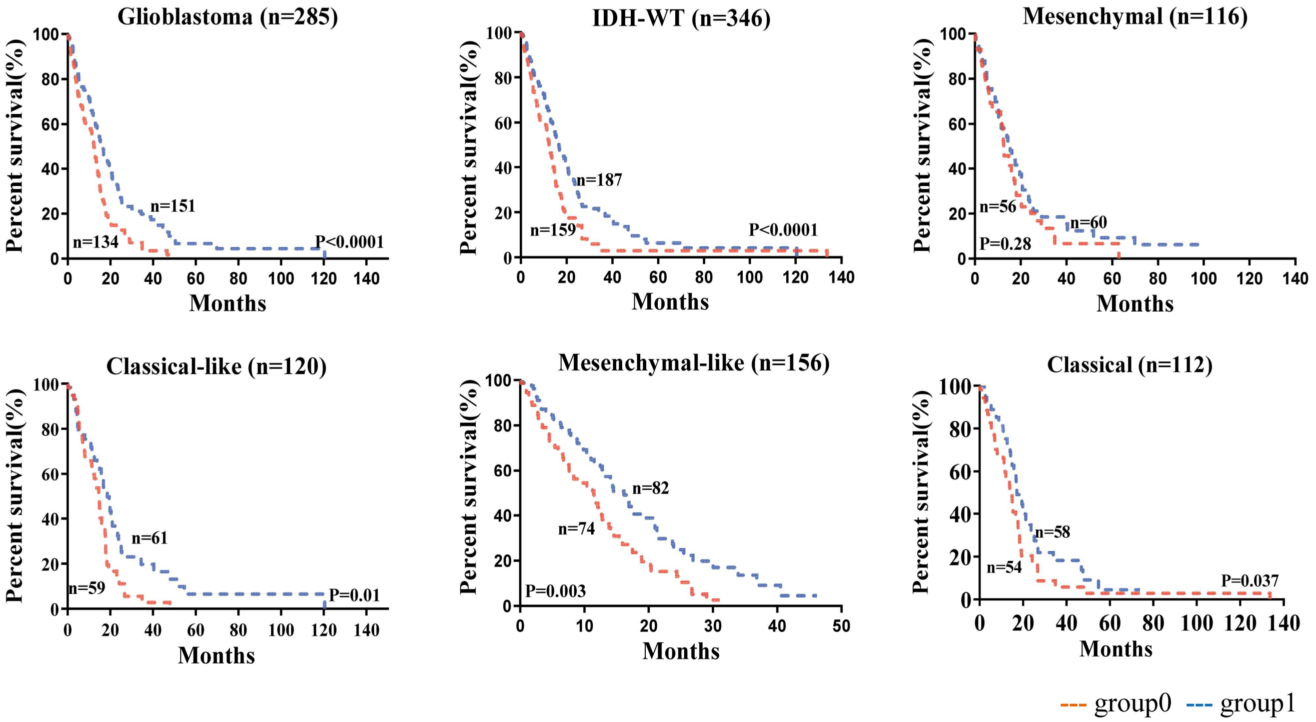


**Supplementary Figure S7**. Survival of TCGA glioma patients stratified by deep learning model .red line, the deep learning model prediction label is 0; blue line, model prediction label is 1. n, number of patients; p-value was determined using the log-rank test.

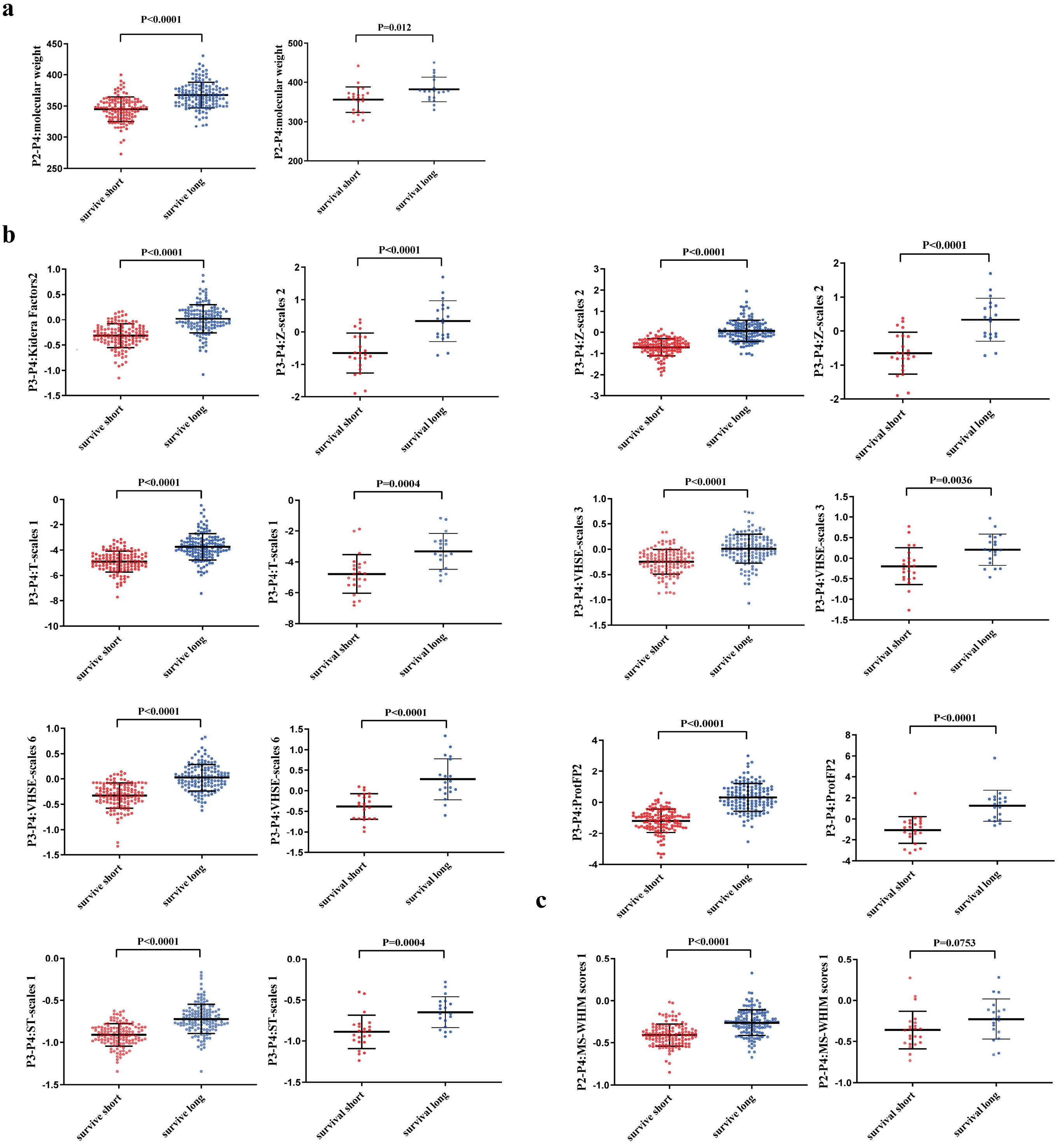


**Supplementary Figure S8**. Comparison of the similarity of valid feature values between long-term survival and short-term survival groups of IDH wild-type GBM in two cohorts. **a**, molecular weight related features. **b**, molecular size/volume related features. **c**, molecular electrostatic potential/polarity related features. P-value was calculated by unpaired T test. ). The left and the right col contain the TCGA cohort and Pri cohort, respectively.

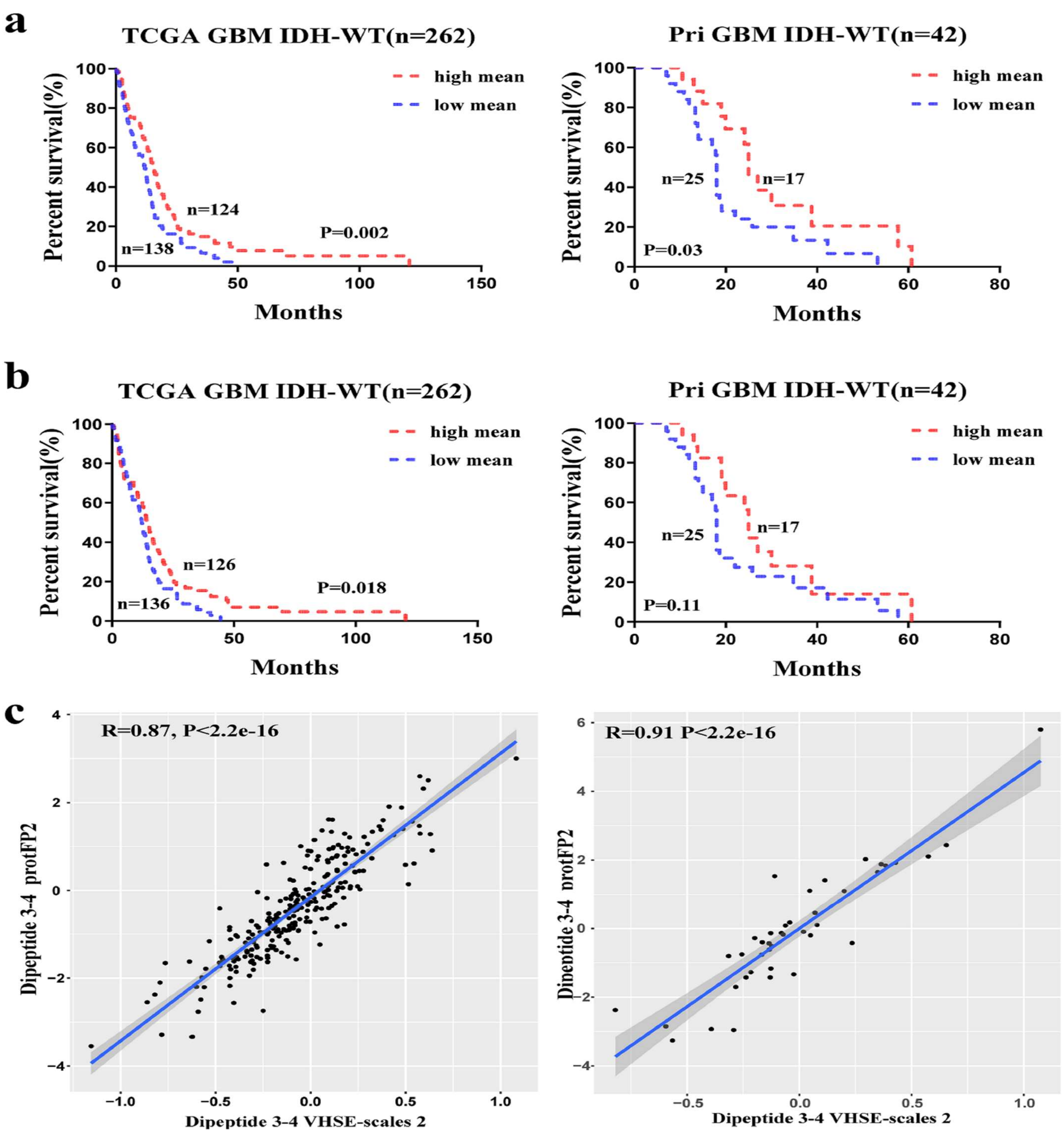


**Supplementary Figure S9.** Survival of glioma patients stratified according to 2 feature values, and analysis of the correlations between these features in two cohorts. **a**, absolute position 3 and 4 composed-dipeptide protFP2 value ; **b**, absolute position 3 and 4 composed-dipeptide VHSE-scale 2. red line, high mean value, blue , low mean value; n, number of patients; p-value was determined using the log-rank test. **c**, the Pearson correlation coefficient of the two features. The left and the right col contain the TCGA cohort and Pri cohort, respectively.

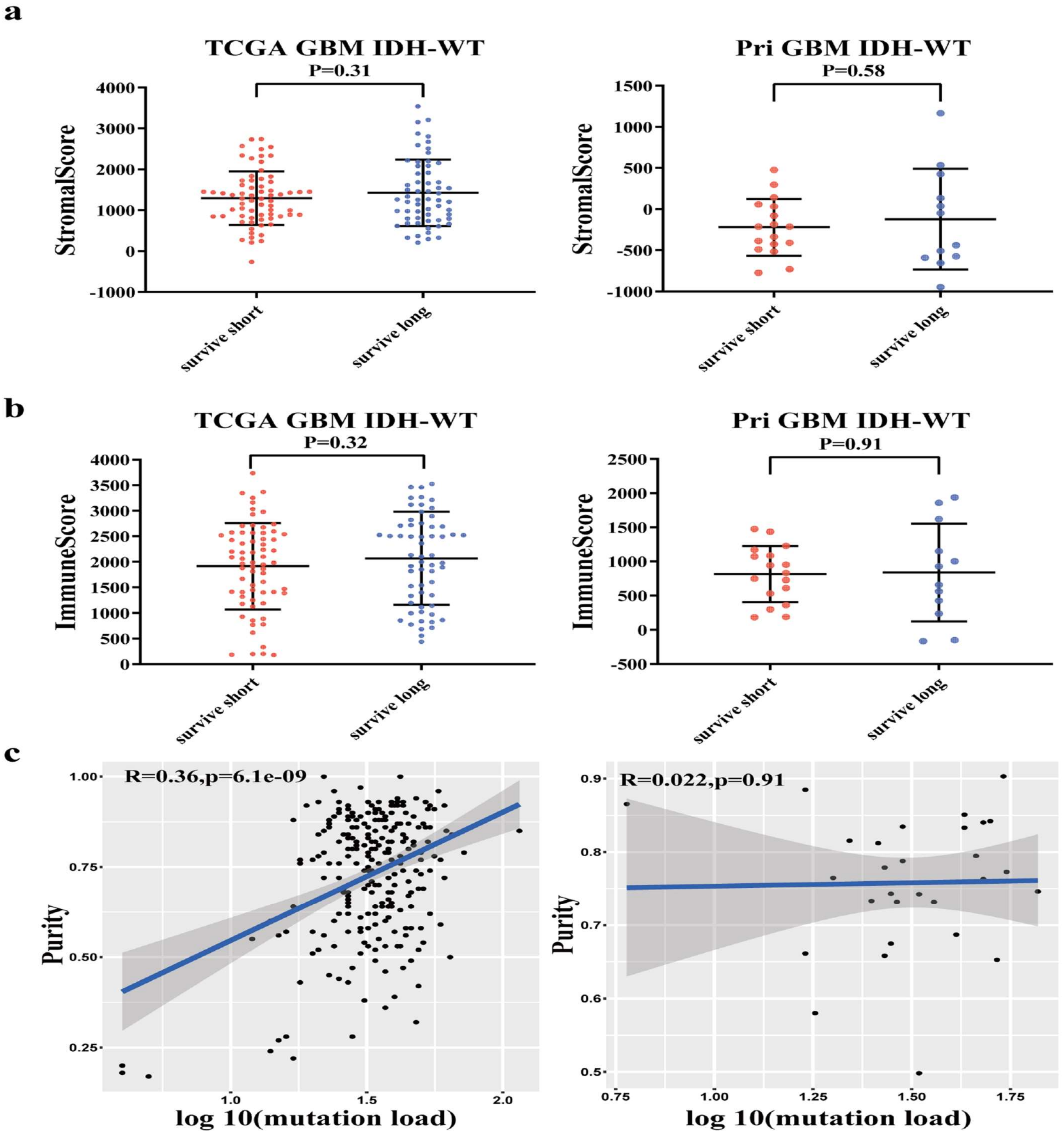


**Supplementary Figure S10**. Comparison of the similarity of immune score and stromal score between two groups and the correlation analysis between purity and mutation load. **a**, stromal score. **b**, immune score. Immune score and stromal score were derived by ESTIMATE. P value was calculated by unpaired T test. **c**, pearson correlation between purity and log10(mutation load). The left and the right col contain the TCGA cohort and Pri cohort, respectively.

**Supplementary Table S1. Multivariate Cox regression analysis including position 3-4 composed-dipeptide VHSE-scale 2 value, mutation load and age for TCGA IDH wild type GBM (n=262).**

| TCGA cohort (n=262) | HR | 95%CI | P |
|---|---|---|---|
| Age | 1.0306 | [1.0163,1.0452] | 2.42e-05 |
| Mutation load | 0.9866 | [0.9729,1.0005] | 0.0595 |
| Dipeptide 3-4 VHSE2 | 0.5732 | [0.3661,0.8972] | 0.0149 |

**Supplementary Table S2. Multivariate Cox regression analysis including position 3-4 composed-dipeptide VHSE-scale 2 value, mutation load and age for Pri IDH wild type GBM (n=42).**

| Pri cohort (n=42) | HR | 95%CI | P |
|---|---|---|---|
| Age | 1.0230 | [0.9914,1.0555] | 0.1555 |
| Mutation load | 1.0047 | [0.9872,1.0226] | 0.6011 |
| Dipeptide 3-4 VHSE2 | 0.3327 | [0.1120,0.9885] | 0.0476 |

**Supplementary Table S3. Multivariate Cox regression analysis including position 3-4 composed-dipeptide protFP 2 value, mutation load and age for TCGA IDH wild type GBM (n=262).**

| TCGA cohort (n=262) | HR | 95%CI | P |
|---|---|---|---|
| Age | 1.0303 | [1.0159,1.0449] | 3.19e-05 |
| Mutation load | 0.9865 | [0.9727,1.0005] | 0.0579 |
| Dipeptide 3-4 protFP2 | 0.8818 | [0.7804,0.9964] | 0.0437 |

**Supplementary Table S4. Multivariate Cox regression analysis including position 3-4 composed-dipeptide protFP 2 value, mutation load and age for Pri IDH wild type GBM (n=42).**

| Pri cohort (n=42) | HR | 95%CI | P |
|---|---|---|---|
| Age | 1.024711 | [0.9936,1.0568] | 0.1208 |
| Mutation load | 1.0052 | [0.9877,1.0230] | 0.5641 |
| Dipeptide 3-4 protFP2 | 0.7868 | [0.6244,0.9915] | 0.0421 |

**Supplementary Table S5. Functional annotation for the lists of genes differentially expressed analyzed by GSEA in TCGA cohort.**

| GENE SET | NAME | SIZE | ES | NES | NOM p-val |
|---|---|---|---|---|---|
| NERVOUS SYSTEM DEVELOPMENT | GO_REGULATION_OF_CELL_SIZE | 161 | -0.3763 | -1.5820 | 0.0200 |
| | GO_CELL_VOLUME_HOMEOSTASIS | 26 | -0.4998 | -1.6130 | 0.0199 |
| | GO_REGULATION_OF_EXTENT_OF_CELL_GROWTH | 98 | -0.3821 | -1.4872 | 0.0432 |
| | GO_NEGATIVE_REGULATION_OF_AXONOGENESIS | 65 | -0.4853 | -1.7992 | 0.0060 |
| | GO_NEGATIVE_REGULATION_OF_CELL_MORPHOGENESIS_INVOLVED_IN_DIFFERENTIATION | 114 | -0.4008 | -1.6149 | 0.0161 |
| | GO_REGULATION_OF_AXONOGENESIS | 163 | -0.3987 | -1.6032 | 0.0236 |
| | GO_NEGATIVE_REGULATION_OF_NEURON_DIFFERENTIATION | 180 | -0.3794 | -1.6456 | 0.0078 |
| | GO_NEGATIVE_REGULATION_OF_NERVOUS_SYSTEM_DEVELOPMENT | 247 | -0.3620 | -1.5871 | 0.0078 |
| | GO_NEGATIVE_REGULATION_OF_CELL_MORPHOGENESIS_INVOLVED_IN_DIFFERENTIATION | 114 | -0.4008 | -1.6149 | 0.0161 |
| | GO_NEGATIVE_REGULATION_OF_CELL_DEVELOPMENT | 283 | -0.3783 | -1.7265 | 0.0000 |
| | GO_REGULATION_OF_GLIAL_CELL_PROLIFERATION | 18 | -0.5012 | -1.5460 | 0.0427 |
| | GO_REGULATION_OF_NEURON_MIGRATION | 27 | -0.5358 | -1.7249 | 0.0059 |
| FOREBRAIN DEVELOPMENT | GO_FOREBRAIN_CELL_MIGRATION | 58 | -0.4223 | -1.6291 | 0.0077 |
| | GO_CEREBRAL_CORTEX_CELL_MIGRATION | 39 | -0.4377 | -1.5670 | 0.0200 |
| | GO_TELENCEPHALON_DEVELOPMENT | 205 | -0.3176 | -1.3907 | 0.0454 |
| | GO_TELENCEPHALON_GLIAL_CELL_MIGRATION | 17 | -0.6088 | -1.7113 | 0.0100 |
| | GO_CEREBRAL_CORTEX_RADIALLY_ORIENTED_CELL_MIGRATION | 25 | -0.5699 | -1.8023 | 0.0078 |
| GLIAL CELL DEVELOPMENT | GO_GLIAL_CELL_MIGRATION | 34 | -0.4837 | -1.7057 | 0.0059 |
| | GO_ASTROCYTE_DEVELOPMENT | 18 | -0.6128 | -1.8593 | 0.0000 |
| | GO_GLIOGENESIS | 169 | -0.3763 | -1.4899 | 0.0264 |
| | GO_GLIAL_CELL_DEVELOPMENT | 73 | -0.4118 | -1.5100 | 0.0336 |
| KIDNEY DEVELOPMENT | GO_UROGENITAL_SYSTEM_DEVELOPMENT | 287 | -0.3096 | -1.4206 | 0.0288 |
| | GO_GLOMERULUS_DEVELOPMENT | 46 | -0.4705 | -1.6512 | 0.0144 |
| | GO_METANEPHRIC_NEPHRON_MORPHOGENESIS | 20 | -0.5068 | -1.5361 | 0.0433 |
| | GO_METANEPHRIC_NEPHRON_DEVELOPMENT | 31 | -0.4970 | -1.6176 | 0.0181 |
| | GO_NEPHRON_DEVELOPMENT | 110 | -0.3656 | -1.5165 | 0.0335 |
| | GO_RENAL_SYSTEM_VASCULATURE_DEVELOPMENT | 18 | -0.6440 | -1.7302 | 0.0161 |
| EPIDERMIS DEVELOPMENT | GO_REGULATION_OF_EPITHELIAL_CELL_DIFFERENTIATION | 112 | -0.3763 | -1.5820 | 0.0200 |
| | GO_NEGATIVE_REGULATION_OF_EPIDERMIS_DEVELOPMENT | 15 | -0.4998 | -1.6130 | 0.0199 |
| | GO_REGULATION_OF_EPIDERMIS_DEVELOPMENT | 59 | -0.3821 | -1.4872 | 0.0432 |
| | GO_REGULATION_OF_EPIDERMAL_CELL_DIFFERENTIATION | 41 | -0.4853 | -1.7992 | 0.0060 |
| | GO_NEGATIVE_REGULATION_OF_EPITHELIAL_CELL_DIFFERENTIATION | 34 | -0.4008 | -1.6149 | 0.0161 |
| DETECTION OF LIGHT STIMULUS | GO_DETECTION_OF_LIGHT_STIMULUS | 52 | -0.3987 | -1.6032 | 0.0236 |
| | GO_DETECTION_OF_VISIBLE_LIGHT | 37 | -0.3794 | -1.6456 | 0.0078 |
| | GO_PHOTOTRANSDUCTION | 39 | -0.3620 | -1.5871 | 0.0078 |
| | GO_PHOTOTRANSDUCTION_VISIBLE_LIGHT | 18 | -0.4008 | -1.6149 | 0.0161 |
| | GO_SKELETAL_MUSCLE_CELL_DIFFERENTIATION | 49 | -0.4770 | -1.6848 | 0.0105 |
| | GO_MUSCLE_TISSUE_DEVELOPMENT | 253 | -0.3172 | -1.4376 | 0.0370 |

| | | | | | |
|---|---|---|---|---|---|
| MUSCLE DEVELOPMENT | GO_CARDIAC_VENTRICLE_MORPHOGENESIS | 62 | -0.4266 | -1.6523 | 0.0102 |
| | GO_VENTRICULAR_CARDIAC_MUSCLE_TISSUE_DEVELOPMENT | 45 | -0.4119 | -1.4828 | 0.0408 |
| | GO_CARDIAC_VENTRICLE_DEVELOPMENT | 103 | -0.3493 | -1.4321 | 0.0459 |
| | GO_CARDIAC_MUSCLE_TISSUE_DEVELOPMENT | 130 | -0.3393 | -1.4517 | 0.0492 |
| | GO_CARDIAC_CHAMBER_MORPHOGENESIS | 103 | -0.3637 | -1.4518 | 0.0424 |
| | GO_ATRIOVENTRICULAR_VALVE_DEVELOPMENT | 19 | -0.5456 | -1.5728 | 0.0269 |
| | GO_ATRIOVENTRICULAR_VALVE_MORPHOGENESIS | 16 | -0.5544 | -1.5179 | 0.0430 |
| | GO_OUTFLOW_TRACT_MORPHOGENESIS | 56 | -0.4390 | -1.6306 | 0.0187 |
| NEGATIVE REGULATION OF ANGIOGENESIS | GO_NEGATIVE_REGULATION_OF_BLOOD_VESSEL_ENDOTHELIAL_CELL_MIGRATION | 24 | -0.3791 | -1.6270 | 0.0103 |
| | GO_NEGATIVE_REGULATION_OF_EPITHELIAL_CELL_MIGRATION | 53 | -0.5702 | -1.5534 | 0.0478 |
| | GO_NEGATIVE_REGULATION_OF_ENDOTHELIAL_CELL_MIGRATION | 39 | -0.3559 | -1.4293 | 0.0400 |
| | GO_NEGATIVE_REGULATION_OF_VASCULATURE_DEVELOPMENT | 79 | -0.4178 | -1.5265 | 0.0346 |
| ENDOTHELIUM DEVELOPMENT | GO_MORPHOGENESIS_OF_AN_ENDOTHELIUM | 16 | -0.5385 | -1.8107 | 0.0021 |
| | GO_ENDOTHELIUM_DEVELOPMENT | 84 | -0.5350 | -1.6806 | 0.0250 |
| | GO_ENDOTHELIAL_CELL_DIFFERENTIATION | 67 | -0.4050 | -1.5413 | 0.0268 |
| | GO_ENDOTHELIAL_CELL_DEVELOPMENT | 43 | -0.4584 | -1.6089 | 0.0335 |
| LIPID KINASE ACTIVITY | GO_REGULATION_OF_PHOSPHOLIPID_METABOLIC_PROCESS | 57 | -0.3955 | -1.5765 | 0.0247 |
| | GO_REGULATION_OF_LIPID_KINASE_ACTIVITY | 45 | -0.6210 | -1.7043 | 0.0185 |
| | GO_REGULATION_OF_PHOSPHATIDYLINOSITOL_3_KINASE_ACTIVITY | 37 | -0.4070 | -1.5577 | 0.0312 |
| | GO_POSITIVE_REGULATION_OF_LIPID_KINASE_ACTIVITY | 30 | -0.4084 | -1.5323 | 0.0271 |
| NEGATIVE REGULATION OF ENDOCYTOSIS | GO_REGULATION_OF_RECEPTOR_INTERNALIZATION | 37 | -0.4643 | -1.5897 | 0.0374 |
| | GO_REGULATION_OF_RECEPTOR_MEDIATED_ENDOCYTOSIS | 76 | -0.4205 | -1.5133 | 0.0452 |
| | GO_NEGATIVE_REGULATION_OF_RECEPTOR_MEDIATED_ENDOCYTOSIS | 17 | -0.4623 | -1.6021 | 0.0245 |
| | GO_NEGATIVE_REGULATION_OF_ENDOCYTOSIS | 38 | -0.5463 | -1.8176 | 0.0082 |
| CELL CYCLE | GO_REGULATION_OF_MEIOTIC_CELL_CYCLE | 37 | -0.5633 | -1.8370 | 0.0041 |
| | GO_REGULATION_OF_REPRODUCTIVE_PROCESS | 119 | -0.4719 | -1.6415 | 0.0144 |
| | GO_POSITIVE_REGULATION_OF_REPRODUCTIVE_PROCESS | 51 | -0.4182 | -1.6087 | 0.0167 |
| | GO_POSITIVE_REGULATION_OF_MULTI_ORGANISM_PROCESS | 148 | -0.6680 | -2.0132 | 0.0022 |
| | GO_REGULATION_OF_MEIOTIC_NUCLEAR_DIVISION | 26 | -0.4526 | -1.5995 | 0.0276 |
| | GO_POSITIVE_REGULATION_OF_CELL_CYCLE | 305 | -0.4041 | -1.5413 | 0.0303 |
| | GO_NEGATIVE_REGULATION_OF_CELL_CYCLE_ARREST | 17 | -0.5100 | -1.8460 | 0.0039 |
| | GO_POSITIVE_REGULATION_OF_CELL_DIVISION | 119 | -0.4054 | -1.4644 | 0.0321 |
| | GO_POSITIVE_REGULATION_OF_NUCLEAR_DIVISION | 59 | -0.6066 | -1.8365 | 0.0060 |
| ACTIN AND CELL BEHAVIOR | GO_ACTOMYOSIN | 53 | -0.4592 | -1.5851 | 0.0148 |
| | GO_POSITIVE_REGULATION_OF_LAMELLIPODIUM_ORGANIZATION | 21 | -0.3330 | -1.4821 | 0.0149 |
| | GO_ACTIN_FILAMENT_BUNDLE | 48 | -0.4802 | -1.7601 | 0.0083 |
| | GO_ACTIN_CYTOSKELETON | 400 | -0.3408 | -1.5084 | 0.0203 |
| | GO_MYOFILAMENT | 22 | -0.5122 | -1.6343 | 0.0191 |
| | GO_ACTIN_BINDING | 362 | -0.3071 | -1.3971 | 0.0472 |
| | GO_FILAMENTOUS_ACTIN | 16 | -0.5672 | -1.6450 | 0.0273 |
| | GO_REGULATION_OF_ACTIN_FILAMENT_DEPOLYMERIZATION | 46 | -0.3676 | -1.6096 | 0.0061 |
| | GO_POSITIVE_REGULATION_OF_PROTEIN_DEPOLYMERIZATION | 18 | -0.4140 | -1.6294 | 0.0121 |
| | GO_POSITIVE_REGULATION_OF_PROTEIN_COMPLEX_DISASSEMBLY | 24 | -0.4813 | -1.6042 | 0.0459 |

**Supplementary Table S6. Functional annotation for the lists of genes differentially expressed analyzed by GSEA in Pri cohort.**

| GENE SET | NAME | SIZE | ES | NES | NOM p-val |
|---|---|---|---|---|---|
| DEVELOPMENT | GO_NEPHRON_EPITHELIUM_DEVELOPMENT | 93 | -0.4751 | -1.6299 | 0.0106 |
| | GO_NEPHRON_DEVELOPMENT | 115 | -0.4694 | -1.5868 | 0.0298 |
| | GO_BRANCHING_INVOLVED_IN_URETERIC_BUD_MORPHOGENESIS | 44 | -0.5077 | -1.6568 | 0.0108 |
| | GO_REGULATION_OF_CELL_PROLIFERATION_INVOLVED_IN_HEART_MORPHOGENESIS | 15 | -0.6520 | -1.6017 | 0.0308 |
| | GO_MESONEPHRIC_TUBULE_MORPHOGENESIS | 53 | -0.4865 | -1.6281 | 0.0168 |
| | GO_METANEPHRIC_NEPHRON_DEVELOPMENT | 32 | -0.5624 | -1.6702 | 0.0261 |
| | GO_MESENCHYMAL_TO_EPITHELIAL_TRANSITION | 15 | -0.5699 | -1.5512 | 0.0306 |
| | GO_REGULATION_OF_MESONEPHROS_DEVELOPMENT | 26 | -0.5620 | -1.6528 | 0.0063 |
| | GO_METANEPHROS_MORPHOGENESIS | 28 | -0.4907 | -1.4784 | 0.0273 |
| | GO_REGULATION_OF_MORPHOGENESIS_OF_A_BRANCHING_STRUCTURE | 53 | -0.5160 | -1.7473 | 0.0021 |
| | GO_KIDNEY_MORPHOGENESIS | 82 | -0.4616 | -1.5773 | 0.0068 |
| | GO_METANEPHROS_DEVELOPMENT | 81 | -0.4828 | -1.6600 | 0.0022 |
| | GO_POSITIVE_REGULATION_OF_KIDNEY_DEVELOPMENT | 41 | -0.4881 | -1.5694 | 0.0107 |
| | GO_POSITIVE_REGULATION_OF_MESONEPHROS_DEVELOPMENT | 22 | -0.5434 | -1.5730 | 0.0145 |
| | GO_RENAL_TUBULE_DEVELOPMENT | 78 | -0.4704 | -1.5806 | 0.0218 |
| | GO_DEVELOPMENTAL_INDUCTION | 27 | -0.5744 | -1.7504 | 0.0022 |
| | GO_MESONEPHROS_DEVELOPMENT | 90 | -0.4135 | -1.4647 | 0.0371 |
| | GO_KIDNEY_EPITHELIUM_DEVELOPMENT | 125 | -0.4409 | -1.5676 | 0.0149 |
| | GO_KIDNEY_MESENCHYME_DEVELOPMENT | 17 | -0.6525 | -1.7126 | 0.0168 |
| | GO_REGULATION_OF_ORGAN_FORMATION | 32 | -0.5590 | -1.8287 | 0.0000 |
| | GO_REGULATION_OF_HEART_MORPHOGENESIS | 29 | -0.5182 | -1.5953 | 0.0165 |
| | GO_ORGAN_INDUCTION | 16 | -0.6573 | -1.8843 | 0.0000 |
| | GO_SMOOTH_MUSCLE_CELL_DIFFERENTIATION | 30 | -0.6063 | -1.6321 | 0.0193 |
| | GO_MUSCLE_CELL_DIFFERENTIATION | 232 | -0.3789 | -1.4548 | 0.0445 |
| | GO_POSITIVE_REGULATION_OF_CARDIAC_MUSCLE_CELL_PROLIFERATION | 19 | -0.5747 | -1.6615 | 0.0143 |
| | GO_MUSCLE_STRUCTURE_DEVELOPMENT | 422 | -0.3797 | -1.4761 | 0.0297 |
| | GO_MYOFILAMENT | 24 | -0.4826 | -1.4807 | 0.0494 |
| | GO_MUSCLE_ORGAN_DEVELOPMENT | 268 | -0.3722 | -1.4283 | 0.0398 |
| | GO_REGULATION_OF_CARDIAC_MUSCLE_CELL_PROLIFERATION | 29 | -0.4985 | -1.5391 | 0.0186 |
| | GO_REGULATION_OF_HEART_GROWTH | 42 | -0.4168 | -1.4310 | 0.0319 |
| | GO_SKELETAL_MUSCLE_ORGAN_DEVELOPMENT | 131 | -0.3886 | -1.4271 | 0.0394 |
| | GO_NEGATIVE_REGULATION_OF_EPITHELIAL_CELL_DIFFERENTIATION | 37 | -0.5377 | -1.7288 | 0.0000 |
| | GO_NEGATIVE_REGULATION_OF_EPIDERMIS_DEVELOPMENT | 16 | -0.5956 | -1.6537 | 0.0063 |
| | GO_REGULATION_OF_EPIDERMIS_DEVELOPMENT | 63 | -0.4213 | -1.4555 | 0.0227 |
| | GO_REGULATION_OF_EPIDERMAL_CELL_DIFFERENTIATION | 45 | -0.5144 | -1.6747 | 0.0082 |
| | GO_EMBRYONIC_SKELETAL_SYSTEM_MORPHOGENESIS | 93 | -0.5174 | -1.6215 | 0.0201 |
| | GO_EMBRYONIC_SKELETAL_SYSTEM_DEVELOPMENT | 121 | -0.4927 | -1.5897 | 0.0308 |
| | GO_EMBRYONIC_FORELIMB_MORPHOGENESIS | 32 | -0.6312 | -1.8401 | 0.0022 |
| | GO_FORELIMB_MORPHOGENESIS | 40 | -0.6591 | -2.0074 | 0.0021 |
| | GO_HINDLIMB_MORPHOGENESIS | 37 | -0.5516 | -1.6244 | 0.0258 |

| | | | | | |
|---|---|---|---|---|---|
| CELL FATE | GO_MUSCLE_CELL_FATE_COMMITMENT | 15 | -0.6617 | -1.7801 | 0.0103 |
| | GO_ECTODERMAL_PLACODE_DEVELOPMENT | 15 | -0.5624 | -1.4811 | 0.0406 |
| | GO_NEURON_FATE_COMMITMENT | 67 | -0.4510 | -1.4405 | 0.0488 |
| | GO_CELL_FATE_COMMITMENT_INVOLVED_IN_FORMATION_OF_PRIMARY_GERM_LAYER | 28 | -0.4790 | -1.4892 | 0.0471 |
| | GO_CELL_FATE_COMMITMENT | 226 | -0.3922 | -1.4485 | 0.0489 |
| DNA TRANSCRIPTION | GO_NUCLEOSOMAL_DNA_BINDING | 30 | -0.5613 | -1.5492 | 0.0398 |
| | GO_ENHANCER_BINDING | 92 | -0.5125 | -1.7069 | 0.0000 |
| | GO_RNA_POLYMERASE_II_DISTAL_ENHANCER_SEQUENCE_SPECIFIC_DNA_BINDING | 64 | -0.5469 | -1.7511 | 0.0020 |
| | GO_TRANSCRIPTION_FACTOR_ACTIVITY_RNA_POLYMERASE_II_DISTAL_ENHANCER_SEQUENCE_SPECIFIC_BINDING | 89 | -0.4780 | -1.5828 | 0.0295 |
| | GO_TRANSCRIPTIONAL_ACTIVATOR_ACTIVITY_RNA_POLYMERASE_II_CORE_PROMOTER_PROXIMAL_REGION_SEQUENCE_SPECIFIC_BINDING | 226 | -0.4546 | -1.6407 | 0.0088 |
| | GO_TRANSCRIPTIONAL_ACTIVATOR_ACTIVITY_RNA_POLYMERASE_II_TRANSCRIPTION_REGULATORY_REGION_SEQUENCE_SPECIFIC_BINDING | 314 | -0.4375 | -1.6303 | 0.0065 |
| | GO_TRANSCRIPTION_FACTOR_ACTIVITY_RNA_POLYMERASE_II_CORE_PROMOTER_PROXIMAL_REGION_SEQUENCE_SPECIFIC_BINDING | 326 | -0.4101 | -1.5091 | 0.0441 |
| | GO_CORE_PROMOTER_PROXIMAL_REGION_DNA_BINDING | 363 | -0.4048 | -1.5120 | 0.0338 |
| | GO_HMG_BOX_DOMAIN_BINDING | 18 | -0.5867 | -1.5328 | 0.0419 |
| CELL DIFFERENCIATION AND PATTERN FORMATION | GO_PROXIMAL_DISTAL_PATTERN_FORMATION | 32 | -0.5980 | -1.8268 | 0.0022 |
| | GO_ANTERIOR_POSTERIOR_PATTERN_SPECIFICATION | 194 | -0.4535 | -1.5828 | 0.0317 |
| | GO_ANTERIOR_POSTERIOR_AXIS_SPECIFICATION | 48 | -0.4958 | -1.5677 | 0.0229 |
| | GO_EMBRYONIC_PATTERN_SPECIFICATION | 58 | -0.4441 | -1.5373 | 0.0185 |
| | GO_AXIS_SPECIFICATION | 90 | -0.4567 | -1.5399 | 0.0235 |
| AA BIOSYNTHETIC AND METABOLIC | GO_METHIONINE_METABOLIC_PROCESS | 18 | -0.6747 | -1.7151 | 0.0040 |
| | GO_SULFUR_AMINO_ACID_BIOSYNTHETIC_PROCESS | 19 | -0.5899 | -1.5468 | 0.0413 |
| | GO_ASPARTATE_FAMILY_AMINO_ACID_BIOSYNTHETIC_PROCESS | 23 | -0.5939 | -1.5312 | 0.0276 |
| | GO_SULFUR_AMINO_ACID_METABOLIC_PROCESS | 40 | -0.5547 | -1.6136 | 0.0079 |
| PHOSPHATIDYLCHOLINE METABOLIC | GO_LYSOPHOSPHOLIPASE_ACTIVITY | 20 | -0.5709 | -1.6967 | 0.0042 |
| | GO_PHOSPHOLIPASE_A2_ACTIVITY | 31 | -0.5016 | -1.6308 | 0.0121 |
| | GO_PHOSPHATIDYLETHANOLAMINE_ACYL_CHAIN_REMODELING | 23 | -0.5646 | -1.6702 | 0.0041 |
| | GO_PHOSPHATIDYLSERINE_METABOLIC_PROCESS | 28 | -0.4944 | -1.5093 | 0.0265 |
| | GO_PHOSPHATIDYLCHOLINE_ACYL_CHAIN_REMODELING | 26 | -0.4962 | -1.5195 | 0.0244 |
| | GO_PHOSPHATIDYLCHOLINE_ACYL_CHAIN_REMODELING | 26 | -0.4962 | -1.5195 | 0.0244 |
| DNA CATABOLIC | GO_APOPTOTIC_DNA_FRAGMENTATION | 15 | -0.6423 | -1.6850 | 0.0084 |
| | GO_DNA_CATABOLIC_PROCESS_ENDONUCLEOLYTIC | 19 | -0.6155 | -1.6531 | 0.0167 |
| | GO_DNA_CATABOLIC_PROCESS | 27 | -0.5160 | -1.4684 | 0.0354 |
| | GO_NUCLEASE_ACTIVITY | 196 | -0.4428 | -1.5150 | 0.0438 |
| | GO_ENDODEOXYRIBONUCLEASE_ACTIVITY | 49 | -0.5508 | -1.7097 | 0.0083 |
| | GO_DEOXYRIBONUCLEASE_ACTIVITY | 65 | -0.5413 | -1.6832 | 0.0042 |
| | GO_EXONUCLEASE_ACTIVITY | 76 | -0.5044 | -1.5316 | 0.0493 |
| | GO_EXODEOXYRIBONUCLEASE_ACTIVITY | 15 | -0.6494 | -1.5721 | 0.0285 |
| | GO_ENDONUCLEASE_ACTIVITY_ACTIVE_WITH_EITHER_RIBO_OR_DEOXYRIBONUCLEIC_ACIDS_AND_PRODUCING_3_PHOSPHOMONOESTERS | 19 | -0.5400 | -1.4735 | 0.0432 |
